# Muller’s ratchet and gene duplication

**DOI:** 10.1101/2024.12.27.630357

**Authors:** Fabian Freund, Johannes Wirtz, Yichen Zheng, Yannick Schäfer, Thomas Wiehe

## Abstract

Copy number of genes in gene families can be highly variable among individuals and may continue to change across generations. Here, we study a model of duplication-selection interaction, which is related to Haigh’s mutation-selection model of Muller’s ratchet. New gene copies are generated by duplication but fitness of individuals decreases as copy number increases. Our model comes in two flavours: duplicates are copied either from a single template or from any existing copy. A duplication-selection equilibrium exists in both cases for infinite size populations and is given by a shifted Poisson or a negative binomial distribution. Unless counteracted by synergistic epistasis, finite populations suffer from loss of low copy-number haplotypes by drift, forcing them into a regime called ‘run-away evolution’ in which new copies accumulate without bound nor equilibrium. We discuss a few empirical examples and interpret them in the light of our models. Generally, large gene families appear too over-dispersed to fit the single template model suggesting a dynamic, and potentially accelerating, duplication process.

## Introduction

In his classical paper (1), Muller studied “the relation of recombination to mutational advance” noting “a kind of irreversible ratchet mechanism in non-recombining species”. The term “Muller’s ratchet” appeared only several years later in the title of Haigh’s paper from 1978, where he studied the dynamics of deleterious mutations in infinite and finite populations (2). While a stable mutation-selection equilibrium is reached in infinite populations void of drift, finite populations continue to accumulate deleterious mutations if not counter-acted by recombination. Together, the two papers triggered broad interest in this fundamental evolutionary mechanism among both theoreticians and experimentalists entailing a large volume of follow-up studies (see (3) for a recent overview). One of the focal questions of theoretical interest concerned the speed of the ratchet. Clearly, this depends on the relative magnitude of the strength of selection and drift and the mutation rate (4–7). Importantly, the ratchet in finite populations can be slowed down and even be halted by appropriate recombination of mutation-loaded chromosomes. This effect keeps the mutation load of natural populations in check and also constitutes the most popular explanation for the evolution and maintenance of sexual reproduction (8–12). Another mechanism capable of preventing build-up of mutation load is a steady influx of beneficial mutations, which may drive the population towards a mutation-selection equilibrium (13). A third way to slow down the speed of the ratchet is by synergistic epistasis: it amplifies the deleterious fitness effect of additional mutations in an already mutation- loaded chromosome (14–16) and poses a selection threshold to unlimited accumulation of deleterious mutations. In this sense, epistasis may be considered a substitute for recombination which should also work under forms of reproduction which are not obligately sexual or diocieous.

In the model considered here, we replace mutation by gene duplication and study gene copy number variation (gCNV) in a population. We consider a hypothetical gene family where new copies are produced by duplication of an existing gene, leading to variable family sizes among individuals. We study two versions of the duplication process: one in which only a template gene may duplicate, called ‘single template model’ (STM), and one in which any copy may duplicate, called ‘compound copy model’ (CCM). Our focus is on the overall count of gene copies in a given individual’s genome and we disregard the effect of intra-chromosomal recombination on this count. For a multiplicative fitness function we derive formulae for the duplication-selection equilibrium in infinite populations. We show that this equilibrium is close to a family of standard distributions, and develop ways of estimating the relevant parameters from data. In simulations we study properties of the two model versions in finite populations and under the effect of epistasis. In particular, we show that the ‘duplication ratchet’ is slowed down, and can come to an effective halt, if synergistic epistasis is sufficiently strong. Finally, we interpret in the light of our model experimentally observed copy number counts of several gene families in human (17) and of the NLR gene family of the vertebrate model species *D. rerio* (18, 19).

## Model

Let a population of constant size *N* ≤ ∞ be propagated in discrete generations *t* = 0, 1, 2, Consider a gene family where each individual of the population has *i*_*o*_ +*i, i* ≥ 0, gene copies and *i*_*o*_ ≥ 1 is the smallest copy number found in this population (*i* can differ between individuals, *i*_*o*_ is identical within the population). In general *i*_*o*_ and *i* are realizations of time-dependent discrete random variables *I*_*o*_(*t*) and *I*(*t*) (again, each individual has its random variable *I*, but there is one population-wide *I*_*o*_). In pangenome terminology, we may think of *I*_*o*_(*t*) and *I*(*t*) representing the core and the accessory parts of this gene family.

Let *y*_*i*_(*t*), *i* ≥ 0, be the relative frequency of (the equivalence class of) individuals with *i*_*o*_ + *i* copies (i.e. those, which have *i* ‘extra’ copies) at time *t*. Clearly, for all *t* ≥ 0 holds Σ _*i*_ ≥ _0_ *y*_*i*_(*t*) = 1. Over time, allele frequencies may change due to duplication, selection or – in the case of finite populations – due to drift. In this latter case, also *I*_*o*_(*t*) may change: it increases once all individuals with exactly *i*_*o*_ copies (i.e., the ‘zero’ class with *i* = 0) are lost.

Further, we assume a fitness function governed by the selection coefficient 0 *< s <* 1 and the epistasis parameter 0 *< β* (cf. 10, 14, 15). Fitness 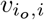 of an individual with *i*_*o*_ + *i* copies is

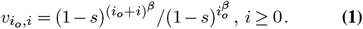

This definition implies that individuals which carry only core genes (*i* = 0), have fitness 1. Mean and variance of fitness in generation *t* are

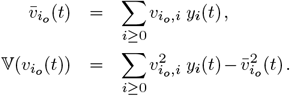

When *β* = 1 there is no epistasis. Fitness is multiplicative and depends only on *i*, but not on *i*_*o*_. In this case

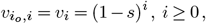

which is identical to the fitness function in Haigh’s model (2, 20). Although our definition of fitness refers to haplotypes, we also use it for genotypes of diploid individuals. Ploidy should be of no concern when assuming random mating and multiplicativity of maternal (*i* copies) and paternal (*j* copies) fitnesses, i.e., 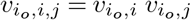. Strictly, this holds only when non-linear fitness effects are neglected and in the absence of epistasis. While the former is unproblematic for small *s*, the effect of epistasis may not easily be hand-waved away and shall be considered in more detail elsewhere. Without epistasis, we have for copy class *i*_*o*_ + *i* after random mating a total frequency 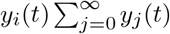 for all genotypes having a class *i* + *i*_*o*_ maternal chromosome. The selection step for these genotypes results in a total frequency

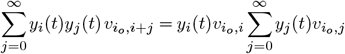

before normalisation. Thus, the selection strength for each maternal class *i* + *i*_*o*_ genotype is proportional to the selection coefficient for the same class in the haploid case. The analogous argument holds for the paternal classes. Consequently, while the normalisation factor may change, haplotype frequencies after selection are the same as in the haploid case. An analogous argument holds if we do consider multiple chromosomes: each chromosome is described by our model, if fitness is multiplicative across the copies on each chromosome.

Consider now the duplication process described by the matrix

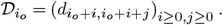

It contains the probabilities 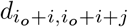 that an individual with *i*_*o*_ + *i* copies has a descendant with *i*_*o*_ + *i* + *j* copies in the next generation. Under duplication, copies are not lost. Hence, *j* ≥ 0. To specify the entries of 𝒟 , we distinguish the single template (STM) and the compound copy (CCM) models. In STM only one gene, the template, can duplicate. Assuming this can happen once per generation, we have

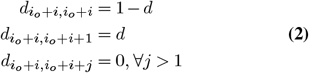

for all *i*_*o*_ *>* 0 and *i* ≥ 0. We call this the Bernoulli model. Allowing that the template may produce more than one copy per generation leads to the Poisson model:

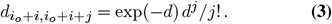

For sufficiently small *d* both Eq. (2) and Eq. (3) are nearly identical. On average *d* new copies are introduced per generation. The variance of the number of new copies is (1− *d*) *d* under the Bernoulli model, and *d* under the Poisson model.

In CCM any copy may duplicate, leading to the binomial probabilities

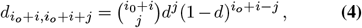

with parameters *i*_*o*_ + *i* and *d*. For *j > i*_*o*_ + *i*, we set 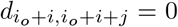, reflecting the assumption that any copy may duplicate at most once per generation. Similarly to STM, this restriction is not imposed under the Poisson approximation, where

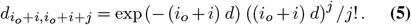

Again, both Eq. (4) and Eq. (5) are nearly identical for small *d*. Both have mean (*i*_*o*_ + *i*) *d* and the variance is (*i*_*o*_ + *i*)(1 − *d*) *d* under the Binomial, and (*i*_*o*_ + *i*) *d* under the Poisson model.

Summarizing, for *N* = ∞ one generation cycle consists of

1. Selection/Normalisation

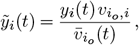
2. Duplication

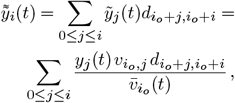
3. Replication

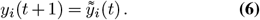

As in classical Wright-Fisher theory, random drift in finite populations (*N* < ∞ ) is realized by multinomial sampling with parameters 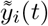 and *N* in step 3. This produces counts, *Y*_*i*_(*t* + 1), of class *i* individuals in generation *t* + 1. The counts are scaled back to relative frequencies by

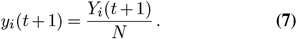

Note, that any class *i* ≥ 0 may be lost by drift if *N* < ∞ Once the class with the currently smallest copy number (*i* = 0) is lost, it cannot be re-surrected by any of the evolutionary forces considered here. Such a loss leads to an increase of *I*_*o*_ from *i*_*o*_ to *i*_õ_ > *i*_*o*_. The indices *i* of all classes are updated to *i −* (*i*_õ_ − *i*_*o*_), such that the new class with 0 extra copies is again labelled with *i* = 0. Each such jump of the random variable *I*_*o*_(*t*) represents a click of the ‘duplication ratchet’. Thus, there is always a non-empty class *i* = 0.

## Results

### Duplication-selection equilibrium

In the absence of drift (*N* = ∞ ) the fittest class (*i* = 0) will not be lost and the count *i*_*o*_ > 0 remains constant over time. Thus, *y*_0_(*t*) is strictly positive for all *t*. Further, without epistasis, a non-trivial duplication-selection equilibrium exists. We present the result for the haploid model with a single chromosome.

#### Proposition 1.

*Let* 0 *< s, d* < 1. *The frequencies* ***y***(*t*) = (*y*_0_(*t*), *y*_1_(*t*),…) *converge in* 𝓁^1^ *to an equilibrium*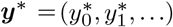, *with recursively determined components*

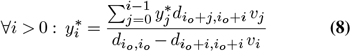

*such that* 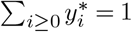

A proof of this proposition is given in the supplementary material. As we have established that copy number development follows the same laws for frequency changes for each haplotype and chromosome, the haploid equilibria and convergence results hold also true in the diploid case, for each haplotype on each chromosome. Thus, to assess the total counts across a diploid genome with *l* chromosomes, we need to sum 2*l* independent random variables, with distribution given by the equilibrium frequencies above, and the core copy numbers for the 2*l* chromosomes.

**STM**. For STM, the recursion Eq. (8) can be solved. With

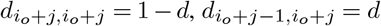 , *d*_*i*_*o*_+*j*−1,*i*_*o*_+*j*_ = *d*, this simplifies to

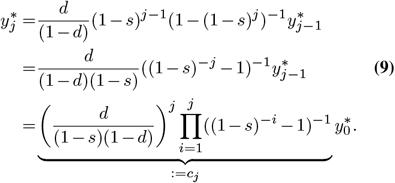

Since 1 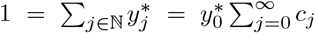, we have 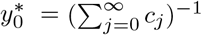 Starting from Eq. (9), we can derive by Taylor approximation an approximate equilibrium distribution for STM:

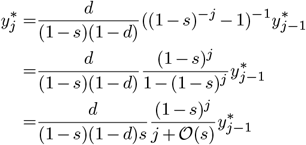

For small *s* and *d*, this becomes approximately

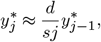

whose unique solution is the Poisson distribution with parameter *d/s*. Thus, the duplication model STM can be approximated by Haigh’s model (2) for the mutation-selection equilibrium for small *s* and *d*. In more detail, at equilibrium, the number of accessory copies *i* is Poisson-distributed with parameter *d/s*, i.e.,

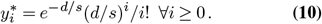

The approximation is very similar to the exact solution, see Table S1. Note the slight difference to Haigh’s model: class 0 refers to those individuals which have 0 accessory copies, but still carry *i*_*o*_ core copies. Thus, mean, variance and skewness of total copy number are

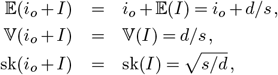

and the *Fano index* (21) (variance to mean ratio) is

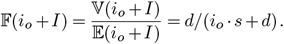

Note that only the mean depends on *i*_*o*_, but neither the variance nor the skewness, because they are invariant under shift by a constant.

An easy calculation shows that equilibrium mean fitness is

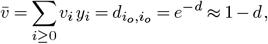

independent of the strength of selection. Fitness variance at equilibrium is

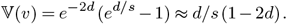

If *d* is small compared to *s*, the fitness variance is approximately *d/s*(1 − 2*d*) ≈ *d/s* and is at the same scale as copy number variance. At equilibrium the effect of selection is balanced by duplication in each generation: after selection and standardization allele frequencies are

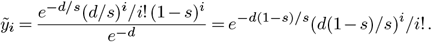

They still follow a Poisson law, but with parameter *d*(1 − *s*)*/s*. I.e., selection reduces mean and variance of copy number by the factor 1 − *s* per generation. This reduction is balanced by duplication. Applying Eq. (3), one has

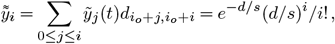

returning allele frequencies to their values before selection and mean copy number to *d/s*. Thus, duplication shifts mean copy number by the factor 1*/*(1 −*s*) ≈ 1 + *s*.

**CCM**. Consider equilibrium mean fitness first. As above, it suffices to satisfy the equilibrium condition for 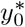

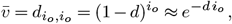

according to Eqs. (4) and (5). The equilibrium condition for class frequencies 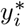 is

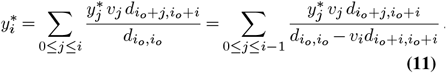

To proceed, consider the ratio 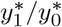. With duplication according to Eq. (5) this becomes

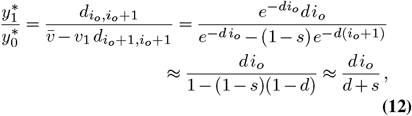

where the last approximation holds if both *s* and *d* are sufficiently small. Similarly, we compute

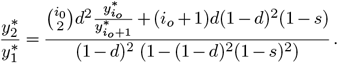

Still, letting *s* and *d* be sufficiently small, and retaining only terms of at most linear order in *s* and *d* in numerator and denominator, we get

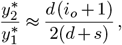

and generally,

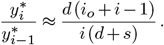

Upon iteration, one finds

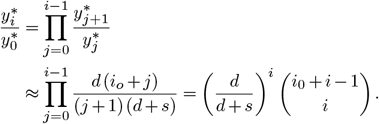

Summation yields

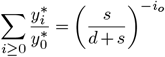

and therefore

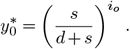

Thus, the equilibrium frequencies of CCM are well approximated by a negative binomial law NB with parameters *i*_*o*_ and *s/*(*d* + *s*) (Table S1). In short,

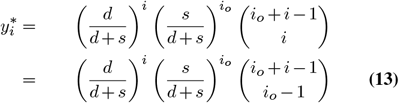

for all *i* ≥ 0. In terminology of the negative binomial distribution 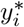 is the probability to draw *i*_*o*_ core copies (‘failures’) before one draws the *i*-th accessory copy (‘success’), given a success probability of *d/*(*d* + *s*). As expected intuitively, the accessory part is large compared to the core part, if *d* is large compared to *s*, and vice versa.

Mean of the accessory part is *i*_*o*_ *d/s*. Mean, variance and skewness of the total copy number at equilibrium are

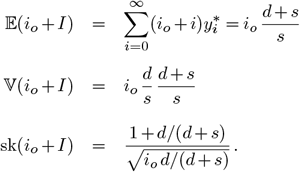

The variance to mean ratio is

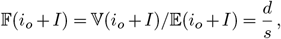

independent of *i*_*o*_, unlike the moments above.

It is worth noting that the random variable *J* = *i*_*o*_ + *I* with *I* ∼ NB(*i*_*o*_, *p*), *p* = *s/*(*s* + *d*) and *p* small, is well approximated by a Gamma-distributed random variable *K* ∼ Γ(*a, b*) with *a* = *i*_*o*_*/*(1 − *p*) = *i*_*o*_(*d* + *s*)*/d* and *b* = *p/*(1 − *p*) = *s/d*. In fact, *K* has the added beauty that it counts the total number of gene copies (which can be directly observed in data) and it is not shifted by a constant. Mean copy number is 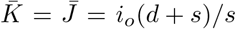 and the Fano index is 𝔽(*K*) = 𝔽(*J*) = *d/s*. Thus, mean and variance of *J* and *K* are identical, but they differ in their skewness: they are 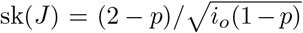 and, slightly smaller,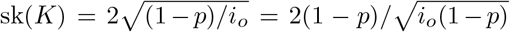. The Gamma-approximation is inaccurate if *p* is not close to 0 or, equivalently, if *s* is large compared to *d*. As the sum of independent Poisson random variables, and also of independent negative binomial distributions and Gamma distributions with equal success probabilities/second parameter are again Poisson/negative binomial/Gamma-distributed random variables, these approximation also hold for the diploid and multi-chromosome case (by adding up the per-chromosome, per-haplotype parameters accordingly).

### Epistasis

Presence of epistasis (*β* ≠1) renders Eqs. (8) and (11) considerably more complicated. Duplication rate and fitness both depend on both variables *i*_*o*_ and *i* (see Eq. (1)). A selection-duplication equilibrium is not readily deduced analytically. Therefore, we resort to simulations (see below) and a qualitative description. Consider the fitness ratio of classes *i* + 1 and *i*

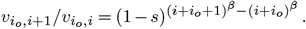

It holds for all *i*_*o*_ ≥ 1

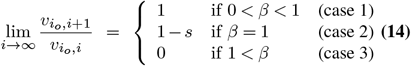

With synergistic epistasis (*β >* 1) and increasing *i*, fitness of class *i* + 1 becomes vanishingly small compared to fitness of class *i*, and thus negative selection against further duplicates becomes increasingly strong. For *β* = 1 the ratio remains constant at 0 < 1− *s* < 1, independent of *i*_*o*_ and *i*. Hence, at equilibrium, we expect 𝔼_*β>*1_(*J*) < 𝔼_*β*=1_(*J*). Since *β* = 1 admits a finite duplication-selection equilibrium in both models STM and CCM, such an equilibrium must also exist for *β* > 1. In contrast, under diminishing epistasis (*β* < 1) the fitness ratio between classes *i* + 1 and *i* tends towards 1 with increasing *i*. Hence, with increasing *i* selection becomes less and less effective to counteract duplication, eventually driving mean copy number to infinity.

The effect of epistasis is more drastic in finite populations, since random drift induces a ratchet effect leading to eventual loss of the least loaded class, i.e. of the class with fewest gene copies, and to a steady increase of *i*_*o*_. Without epistasis (*β* = 1) this happens at constant speed in the STM model. In contrast, in the CCM model the ratchet accelerates with increasing copy number due to the increasing duplication rate. This acceleration can - to some extent - be counteracted by synergistic epistasis (*β* > 1). On the other hand, diminishing returns (*β* < 1) always exacerbates the acceleration.

### Finite population size

The discrete Markov process ***Y*** (*t*) = (*Y*_*i*_(*t*))_*i* ≥ 0_ is governed by a multinomial transition law

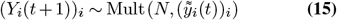

with parameters *N* < ∞ and 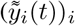 defined in Eq. (6). In the course of one generation frequencies (*Y*_*i*_(*t*))_*i*_ change by duplication, selection and multinomial sampling. In every generation *t* there is a non-zero probability that the least loaded class, i.e. the class *I*_*o*_ with the fewest number of gene copies, is lost by drift. In other words, the random variable *I*_*o*_(*t*) increases over time in a ratchet-like process. Let *T*_*i*_ be a hitting time: the first time 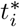 when *I*_*o*_(*t*) = *i*. This means that from 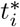 onwards all individuals carry at least *i* copies . We call *R*_*i*_ = 𝔼(*T*_*i*_ − *T*_*i*−1_) the (average) *interarrival time* between consecutive ‘clicks’. The *speed* of the ratchet is the inverse of the interarrival time.

In Haigh’s model (2) this speed is constant and does not depend on *i* (except for the time to the first click, which depends on initial conditions). Haigh had derived an approximation and pointed out that the size of the currently fittest class is the major determinant of the interarrival time. However, outside of a narrow parameter range, his approximation is quite imprecise. Essentially, the problem remains still open despite several attempts of improved approximations (e.g., (4, 6)).

Here, we investigated interarrival times with computer simulations and compared the effect of epistasis in STM and CCM models. We traced the stochastic process (**Y**(*t*))_*t*_ starting from the initial distribution **Y**(0) with *Y*_0_(0) = *N* and *Y*_*i*_(0) = 0 ∀ *i* > 0.

We terminate a simulation run at *T*_2000_, or when *t* = 50 million generations are reached, whatever comes first. Given the equivalence of STM (without epistasis) with Haigh’s model, it is not surprising that the interarrival times are constant in this case (Fig 1A). In contrast, under CCM *R*_*i*_ quickly decrease with increasing *i* (Fig 1B), reflecting the increasing effective duplication rate when the number of copies grows. Besides their strong dependence on the relative magnitudes of *d* and *s*, interarrival times increase with increasing population size.

**Figure 1.**
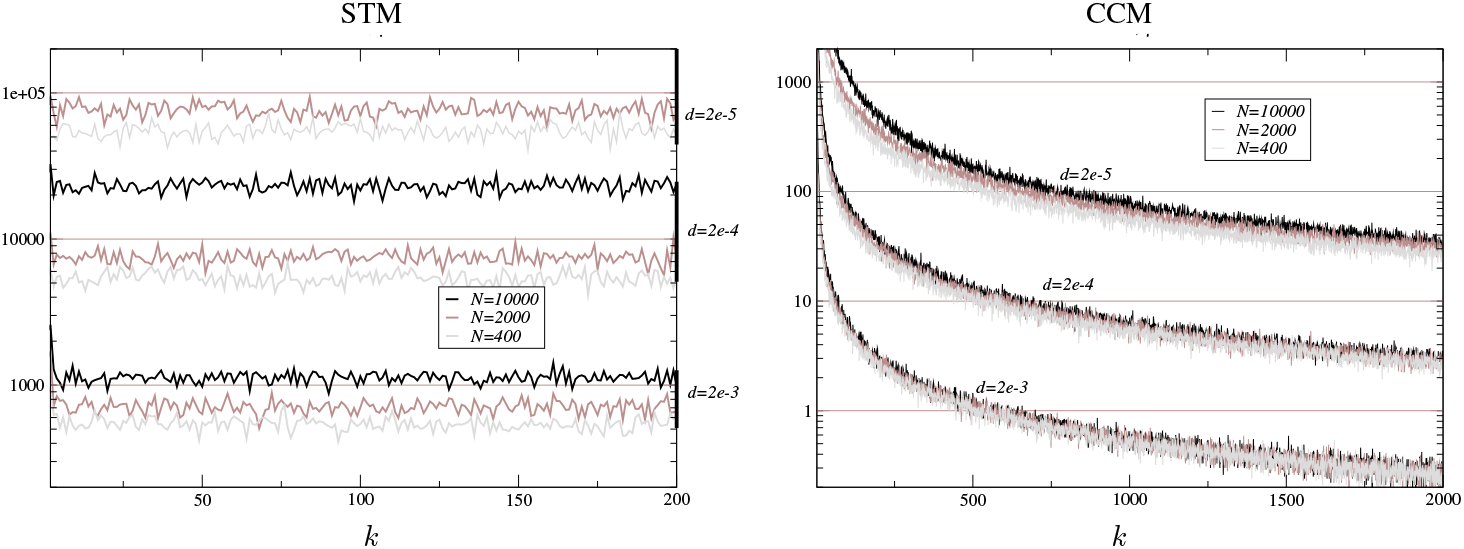
Mean interarrival times *R*_*k*_ (across 100 iterations) *vs*. ratchet ‘clicks’ *k*. Without epistasis (*β* = 1). Simulation results for STM (left) and CCM-models (right). For all cases: *s* = 2 · 10^−4^. Duplication rate from *d* = 2 · 10^−5^ to *d* = 2 · 10^−3^, population size from *N* = 400 to *N* = 10, 000 (see legend in panels). Case STM and *N* = 10, 000 is not shown because interarrival times were exceedingly long in our simulation.

In both STM and CCM the ratchet is slowed down under synergistic (*β* > 1) epistasis. If sufficiently strong, the ratchet can even come to a stop. Operationally, this means that mean copy number 𝔼(*J*(*t*)) remains bounded on a time scale of *O*(10^6^). To be conservative, we ran extra simulations for 10M generations, recorded the population status every 20 generations, and explored the effect of epistasis (*β* ∈ [1.5, 2.1]) on slowing the ratchet (Figure S1). For instance, for *β* = 2 the ratchet came to a stop on the time scale of our simulation with values of 𝔼(*J*) from 1.12 (for *s* = 0.001, *d* = 0.0001, strong selection weak duplication, *d/s* = 0.1) to 49.71 (for *s* = 0.0002, *d* = 0.0005, weak selection strong duplication, *d/s* = 2.5). In contrast, epistasis with *β* = 1.5 was not sufficient to halt the ratchet. Even with *β* = 1.7 duplications accumulate and *T*_1000_ is quickly reached. However, with *β* ≥ 1.8 a slow-down is observed at about 0.3-0.5M generations, and the ratchet comes effectively to a halt at about 2-4M (see Fig 2). The threshold value of *β* between runaway expansion and stopped ratchet generally lies between 1.7 and 1.8 but depends on the value of *s, d* and *N* . The relationship between these parameters is potentially highly complex and non-linear, and is thus beyond the scope of this study.

**Figure 2.**
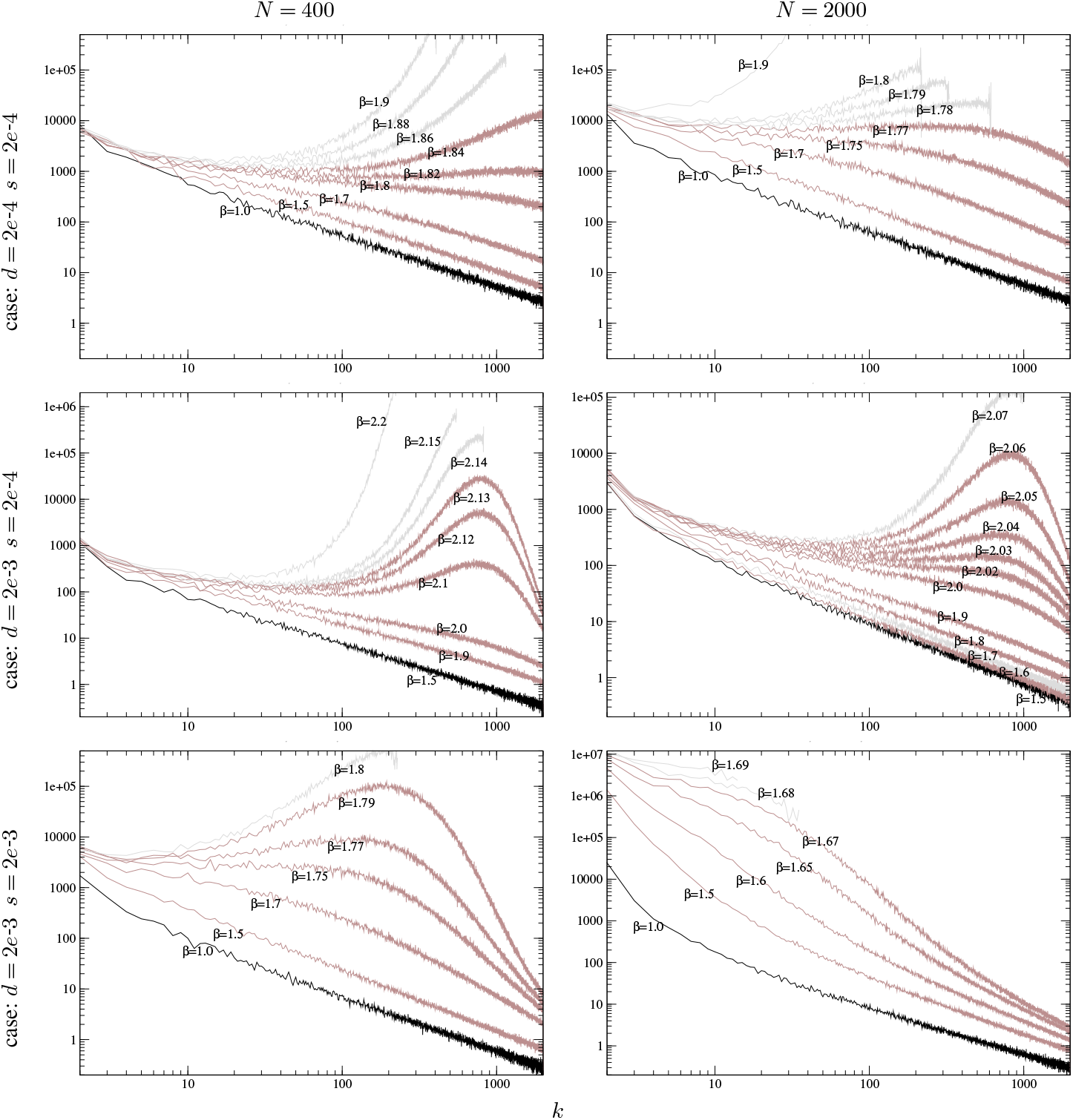
Mean interarrival times *R*_*k*_ *vs*. ratchet ‘clicks’ *k*. CCM model with epistasis (*β* ≥ 1). Mean interarrval times calculated across 100 iterations unless the max number of generations (2*·*10^7^) was exceeded. However, in all cases a minimum number of 10 iterations was taken to calculate the mean. This is why some trajectories terminate before 2000 clicks were reached. Left: *N* = 400. Right: *N* = 2000. Top: *d* = *s* = 2*e*-4, middle: *d* = 2*e*-3 *s* = 2*e*-4, bottom: *d* = *s* = 2*e*-3. Initial configuration for all simulation runs: *Y*_1_ (0) = *N, Y*_*i*_(0) = 0 ∀*i* > 0.

Interestingly, a regime of synergistic epistasis may lead in CCM to a transient behavior of the ratchet: initial slow down, but ultimate acceleration (Fig 2). This suggests the existence of a copy number threshold beyond which the dynamics enters irreversibly a regime of what can be called ‘run-away evolution’.

### Parameter estimation and model selection

Given a sample of *n* counts ***J*** = (*J*_1_, *J*_2_,…, *J*_*n*_) = (*I*_*o*_ + *I*_1_, *I*_*o*_ + *I*_2_,…, *I*_*o*_ + *I*_*n*_) one can define the following estimators for the parameters of the Poisson distribution associated to STM (with parameter *λ*) and of the negative binomial distribution assocuated to CCM (with parameters *i*_*o*_ and *p* = *s/*(*d* + *s*))). Let 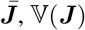 and 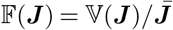 be sample mean, variance and dispersion, respectively, and

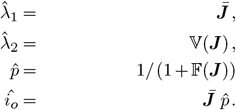

Note that the relationship 𝔽(***J*** ) = (1 − *p*)*/p* = *d/s* yields an estimate of the ratio *d/s* for CCM.

Plausibility of STM may easily be checked by asking whether 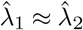. Also, it holds that an under-dispersed negative binomial (i.e., *p* < 1) converges to a Poisson distribution with parameter *λ* = *i*_*o*_ *d/s* when letting *p*↑ 1 and keeping the mean *i*_*o*_ *d/s* of the negative binomial constant. If ***J*** were (nearly) Poisson-distributed with parameter *i*_*o*_ *d/s*, one still would expect its dispersion to be at most one. Thus, observing a dispersion larger than one is incompatible with a STM model. As mentioned above, the (compound) random variable *J* = *I*_*o*_ + *I* with *I*_*o*_ = *i*_*o*_ and *I* ∼ NB(*i*_*o*_, *p*) is well approximated by a Gamma-distributed variable *K* with *K* ∼ Γ(*a, b*) = Γ(*i*_*o*_(1 + *s/d*), *s/d*). It is straightforward to estimate parameters *a, b* from sample ***J*** . We have

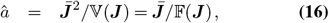

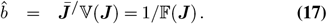

By defintion, 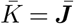 and 𝔽(*K*) = 𝔽(***J*** ).

An alternative way to both distinguish between CCM and STM models and estimate parameters *d/s* and *i*_*o*_ is by maximum-likelihood grid search. Given a sample of gene copy counts, we compute the log-likelihood of the sample for *i*_*o*_ ∈ {1, 2,…, *m* }, where *m* is the minimum of the counts in the sample and with *d/s* ∈ {1*/*100, 1*/*99.75, 1*/*99.5,… , 1*/*1.25, 1, 1.25, 1.5, 1.75,…, 99.75, 100 }. For distinguishing between CCM and STM, we adopt the Kass-Raftery scale interpretation of likelihood quotients (as these are technically Bayes factors). To be precise, if the log-likelihood is > 1 (< −1), we have positive evidence for a better fit of CCM (STM). A log-likelihood above 3 (below -3), will be interpreted as strong evidence for the favoured model. Seen as Bayesian model selection with equal priors for STM and CCM, this means that the posterior probability of the other hypothesis is at most 0.05. In that same line of thought, we can also control that the ‘family-wise error’, i.e. the summed probabilities of the ‘other’ hypotheses, across multiple comparisons to be below some (significance) threshold (which we will set at 0.05). See Section Suppl 4 for details.

### Application I: NLR genes in *Danio rerio*

Schäfer et al. (19) determined copy numbers of the huge NLR gene family in two lab and four wild populations of zebrafish (*Danio rerio*). NLR genes are divided into groups, characterized by certain protein domain signatures. The so-called ‘Fisnacht’ domain is characteristic of all zebrafish NLR genes, while the ‘B30.2’ domain occurs only in some of the groups. We reproduce the gene counts, based on capture of either of the domains, in Supplementary Table S2. Summary statistics of mean, variance and dispersion, together with the estimates of *d/s* and *i*_*o*_, both from moment estimates and from maximum likelihood estimation, are shown in Table 2 below. The parameters *a* and *b* are estimated from the data as described above.

**Table 1.**
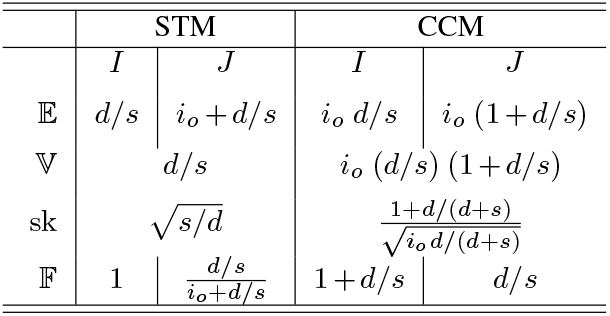
Mean 𝔼, Variance 𝕍 and dispersion 𝔽 = 𝕍*/*𝔼 for STM and CCM of random variables *I* (accessory copy number) and *J* (total copy number) in terms of parameters *i*_*o*_, *d* and *s*. Note that *J* is expected to be under-dispersed (*F* < 1) in STM.

**Table 2.**
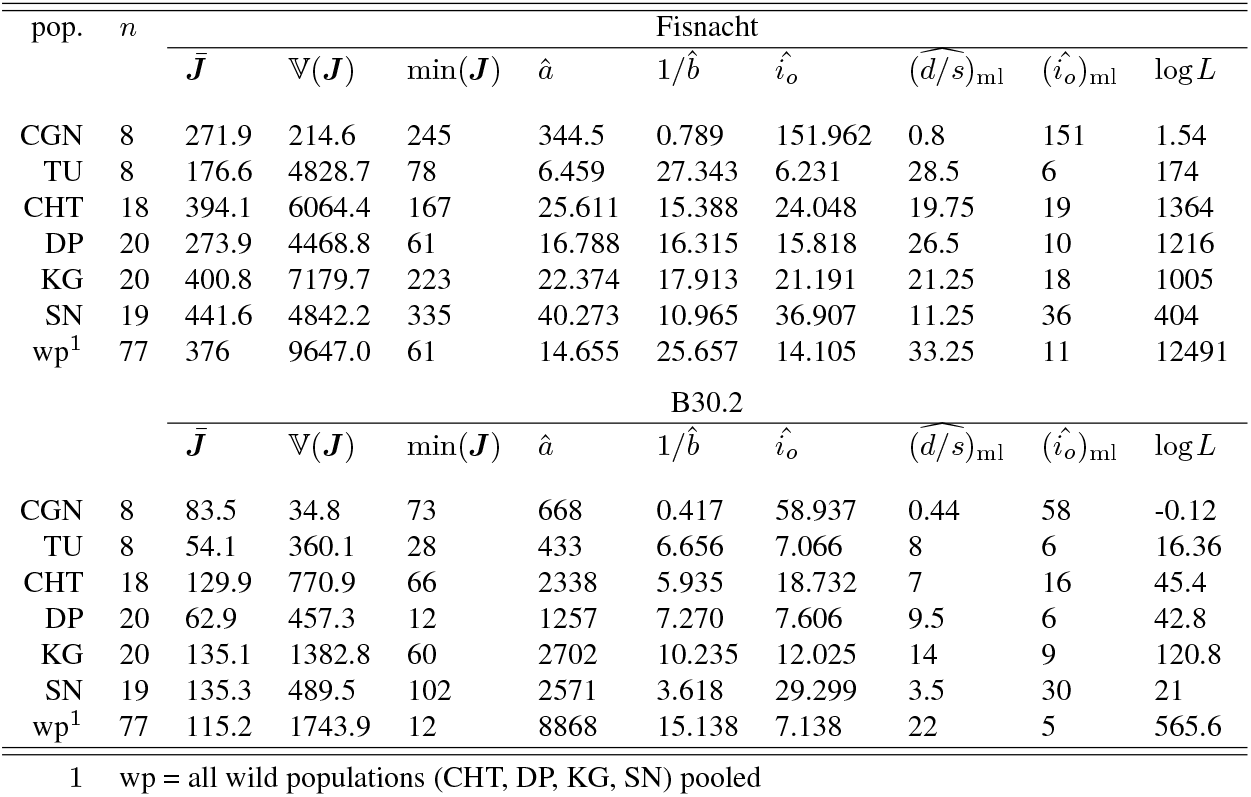
Summary statistics calculated from counts of NLR genes in four populations of *Danio rerio*. Lab strains: CGN and TU; wild populations: CHT, DP, KG and SN. Column *n* refers to the number of individuals which were typed. Further columns: mean, variance and minimum copy number observed. Estimates of parameters *a* and *b* of a Gamma distribution as given in Eqs (16) and (17). Core copy number 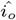 is estimated as 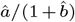. Note that 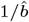 is an estimator of the ratio *d/s*. Moreover, we report the maximum likelihood estimates of *d/s* and *i*_*o*_ for CCM, as well as the log likelihood of CCM over STM.

| pop. | $n$ | Fisnacht | | | | | | | | |
| --- | --- | --- | --- | --- | --- | --- | --- | --- | --- | --- |
| | | $\bar{J}$ | $\mathbb{V}(J)$ | $\min(J)$ | $\hat{a}$ | $1/\hat{b}$ | $\hat{i}_o$ | $(\widehat{d/s})_{\text{ml}}$ | $(\hat{i}_o)_{\text{ml}}$ | $\log L$ |
| CGN | 8 | 271.9 | 214.6 | 245 | 344.5 | 0.789 | 151.962 | 0.8 | 151 | 1.54 |
| TU | 8 | 176.6 | 4828.7 | 78 | 6.459 | 27.343 | 6.231 | 28.5 | 6 | 174 |
| CHT | 18 | 394.1 | 6064.4 | 167 | 25.611 | 15.388 | 24.048 | 19.75 | 19 | 1364 |
| DP | 20 | 273.9 | 4468.8 | 61 | 16.788 | 16.315 | 15.818 | 26.5 | 10 | 1216 |
| KG | 20 | 400.8 | 7179.7 | 223 | 22.374 | 17.913 | 21.191 | 21.25 | 18 | 1005 |
| SN | 19 | 441.6 | 4842.2 | 335 | 40.273 | 10.965 | 36.907 | 11.25 | 36 | 404 |
| wp<sup>1</sup> | 77 | 376 | 9647.0 | 61 | 14.655 | 25.657 | 14.105 | 33.25 | 11 | 12491 |
| B30.2 |  |  |  |  |  |  |  |  |  |  |
| pop. | $n$ | $\bar{J}$ | $\mathbb{V}(J)$ | $\min(J)$ | $\hat{a}$ | $1/\hat{b}$ | $\hat{i}_o$ | $(\widehat{d/s})_{\text{ml}}$ | $(\hat{i}_o)_{\text{ml}}$ | $\log L$ |
| | | $\bar{J}$ | $\mathbb{V}(J)$ | $\min(J)$ | $\hat{a}$ | $1/\hat{b}$ | $\hat{i}_o$ | $(\widehat{d/s})_{\text{ml}}$ | $(\hat{i}_o)_{\text{ml}}$ | $\log L$ |
| CGN | 8 | 83.5 | 34.8 | 73 | 668 | 0.417 | 58.937 | 0.44 | 58 | -0.12 |
| TU | 8 | 54.1 | 360.1 | 28 | 433 | 6.656 | 7.066 | 8 | 6 | 16.36 |
| CHT | 18 | 129.9 | 770.9 | 66 | 2338 | 5.935 | 18.732 | 7 | 16 | 45.4 |
| DP | 20 | 62.9 | 457.3 | 12 | 1257 | 7.270 | 7.606 | 9.5 | 6 | 42.8 |
| KG | 20 | 135.1 | 1382.8 | 60 | 2702 | 10.235 | 12.025 | 14 | 9 | 120.8 |
| SN | 19 | 135.3 | 489.5 | 102 | 2571 | 3.618 | 29.299 | 3.5 | 30 | 21 |
| wp<sup>1</sup> | 77 | 115.2 | 1743.9 | 12 | 8868 | 15.138 | 7.138 | 22 | 5 | 565.6 |

1 wp = all wild populations (CHT, DP, KG, SN) pooled

Except for population CGN, we observe over-dispersed NLR copy counts. In most populations the Fano index is substantially larger than one. This observation is compatible with CCM, but not with STM. This is in agreement with the maximum likelihood grid search, where we find no evidence for STM (all log likelihood ratios bigger than -1) for either gene family, but at least positive evidence for all population but one (log likelihood ratio bigger than 1) and strong evidence for all but two (log likelihood ratio bigger than 3) population across both gene families (Table 2), even when controlling across all tests simultaneously (log likelihood threshold < 5.64, see Section Suppl 4). Moreover, moment and maximum likelihood estimates agree very well (Table 2) and there is an overall good fit of the CCM model to the data (Figure 3) yielding reliable parameter estimates. The discrepancy in dispersion between the lab population CGN and the other populations is most likely to be attributed to population history. CGN is a heavily inbred population of small effective size, established more recently than, and derived from, the other lab population in our sample (TU). This could have led to a strong founder effect due to a small number of founding individuals with a particularly large copy number.

**Figure 3.**
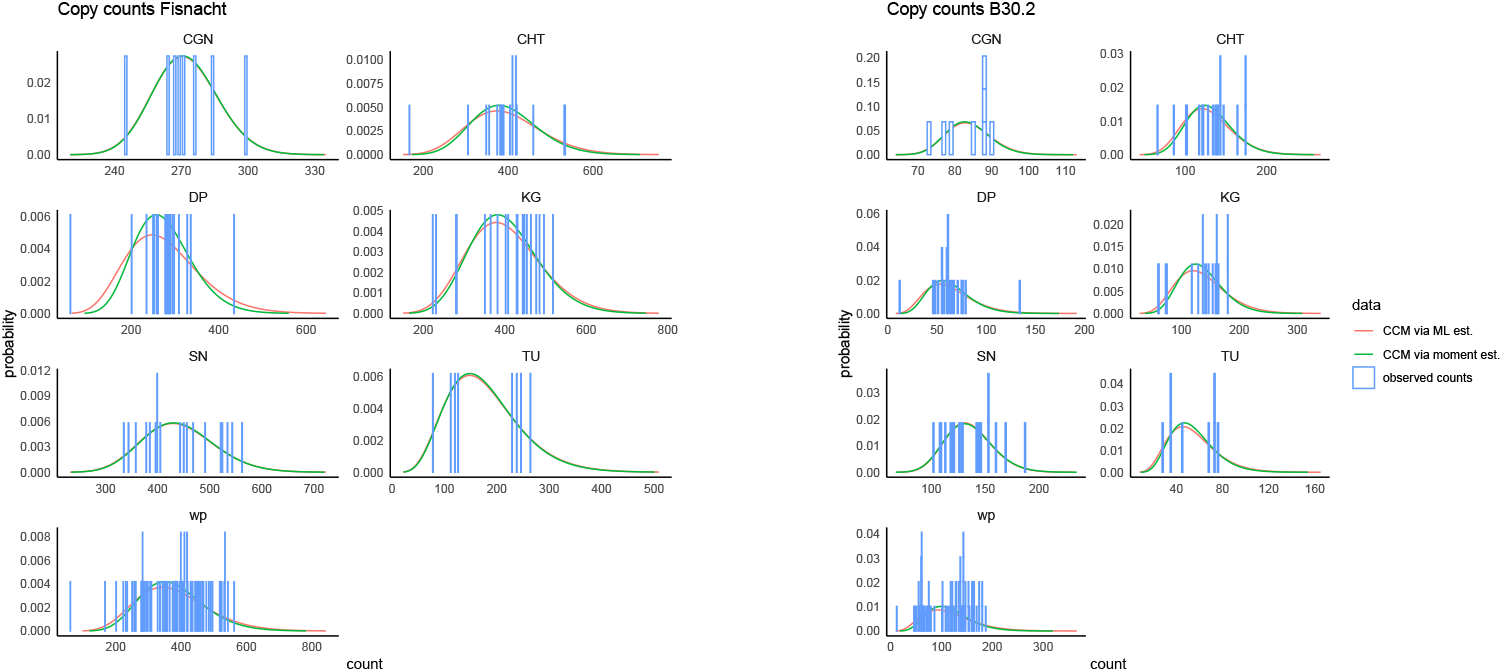
Empirical gene copy count distribution (blue bars) and lineplots of the probabilities of counts for the fitted CCM distribution (using maximum likelihood or momentbased parameter estimators), for two gene domains (Fisnacht, B30.2) from samples of lab (CGN,TU) and wild populations (CHT,DP,KG,SN) of *D. rerio*. wp: wild populations pooled. The observed counts are listed in Table S2.

**Figure 4.**
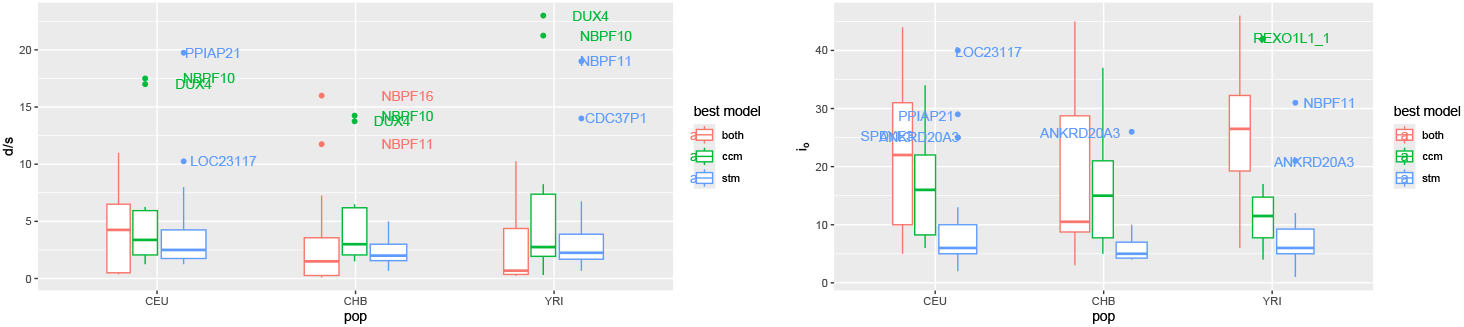
Left: boxplot of *d/s* estimates (log-scaled ordinate). Right: boxplot of *i*_*o*_ estimates in 52 human gene families and three populations, listed in Table 4.

### Application II: Gene families in three populations of human

Brahmachary et al. (17) reported a survey of copy number variation in about 150 gene families (Suppl Table S3) in the three human populations YRI, CEU and CHB. Excluding loci with little variation (less than five copies in YRI, less than two copies in any of the populations), genes on sex chromosomes, and microsatellites, we selected a subset of 52 gene families (Table 4) for which we determined sample statistics and used our maximum likelihood approach to perform model selection and to estimate core copy number *i*_*o*_ and the ratio *d/s* under the better fitting model from CCM and STM (as explained above). The gene families tend to fit better to the STM model than the CCM model in all three populations, albeit 19-38% of gene families not clearly supporting one model over the other (Table 3). While gene families do not always fit to the same model in each population, the model preference, measured as the log likelihood ratio, is strongly correlated between families (Spearman/Pearson correlations of 0.59/0.98 for YRI:CEU, 0.64/1 for YRI:CHB, 0.79/0.99 for CEU:CHB; we report additionally Spearman rank correlation as the log likelihood ratios spread unevenly across their range). Most human gene families we analysed tend to average below 50 copies, thus clearly fewer copies than the NLR genes in *D. rerio*. There is a positive correlation between average copy number and log likelihood ratios (Figure 5): Across populations, Pearson correlations are at least 0.89, while rank (Spearman) correlations are at least 0.67. This indicates a tendency of larger families to fit CCM better. In particular, the largest gene families with more than 100 copies (DUX4 (22), USP17, NBPF10, REXO1L1) all have strong support for the CCM model (log BF > 8.05, the threshold for strong evidence corrected for multiple testing), as shown in Figure 5.

**Table 3.** Proportion of selected model (CCM, STM or no positive evidence for either) of gene families in three human populations (YRI, CEU and CHB; rounded to two digits). We consider a Bayes factor with | log(*BF* )| > 1 positive evidence.

| Population | CCM | STM | no evidence |
| --- | --- | --- | --- |
| YRI | 0.19 | 0.62 | 0.19 |
| CEU | 0.19 | 0.63 | 0.17 |
| CHB | 0.19 | 0.42 | 0.38 |

**Table 4.**
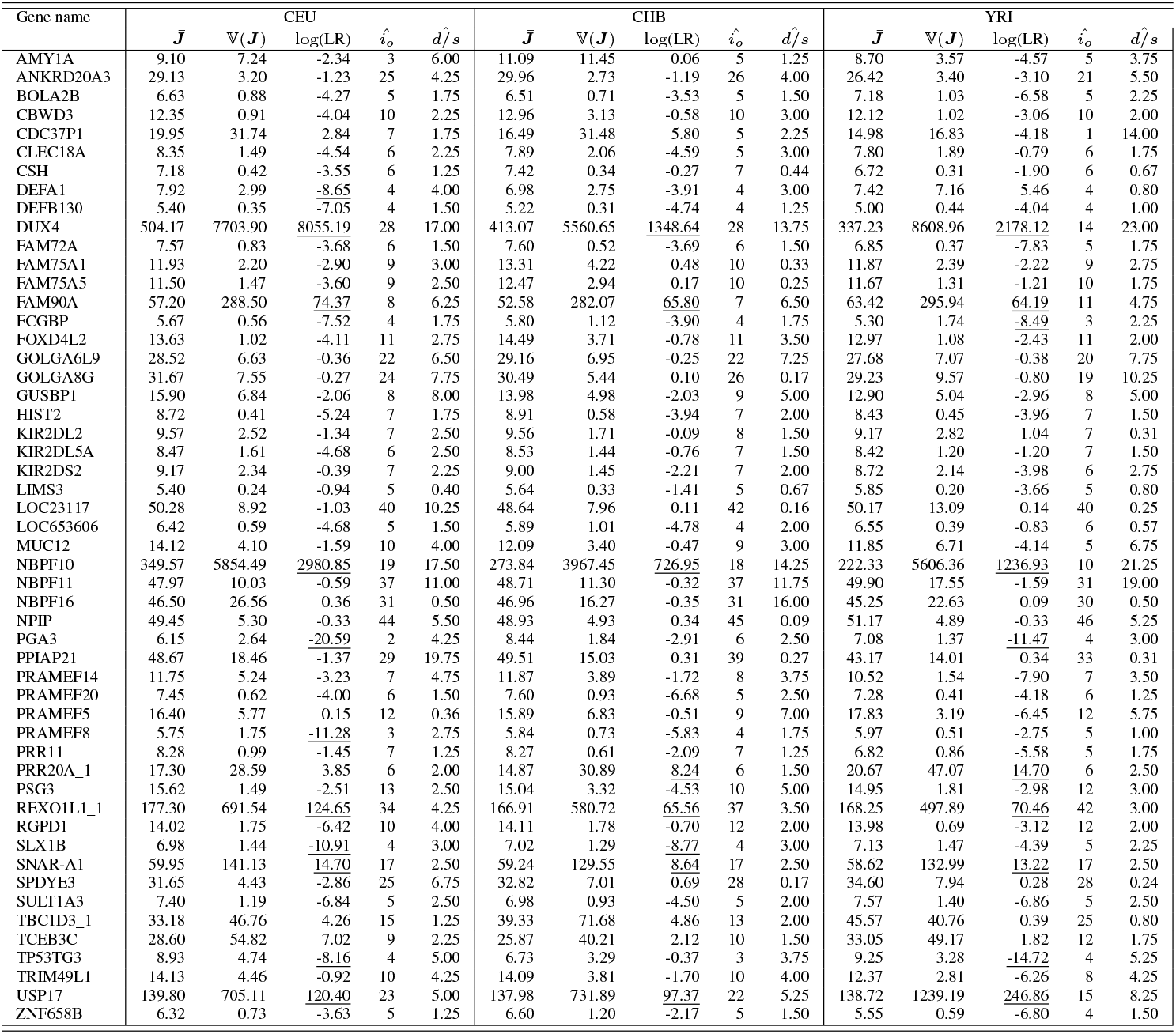
Gene families in three human populations (CEU, CHB and YRI). Data from (17) and adapted by (23). Columns: mean 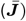 and variance (𝕍(***J*** ) of observed copy number. Log likelihood ratios (log(LR)) between CCM and STM, and maximum likelihood estimates of *i*_*o*_ and *d/s*. Underlined log(LR) denote gene families with strong support after correction for all (3×52) comparisons in this table (see Sect. Suppl 4, threshold *±*8.05)

**Figure 5.**
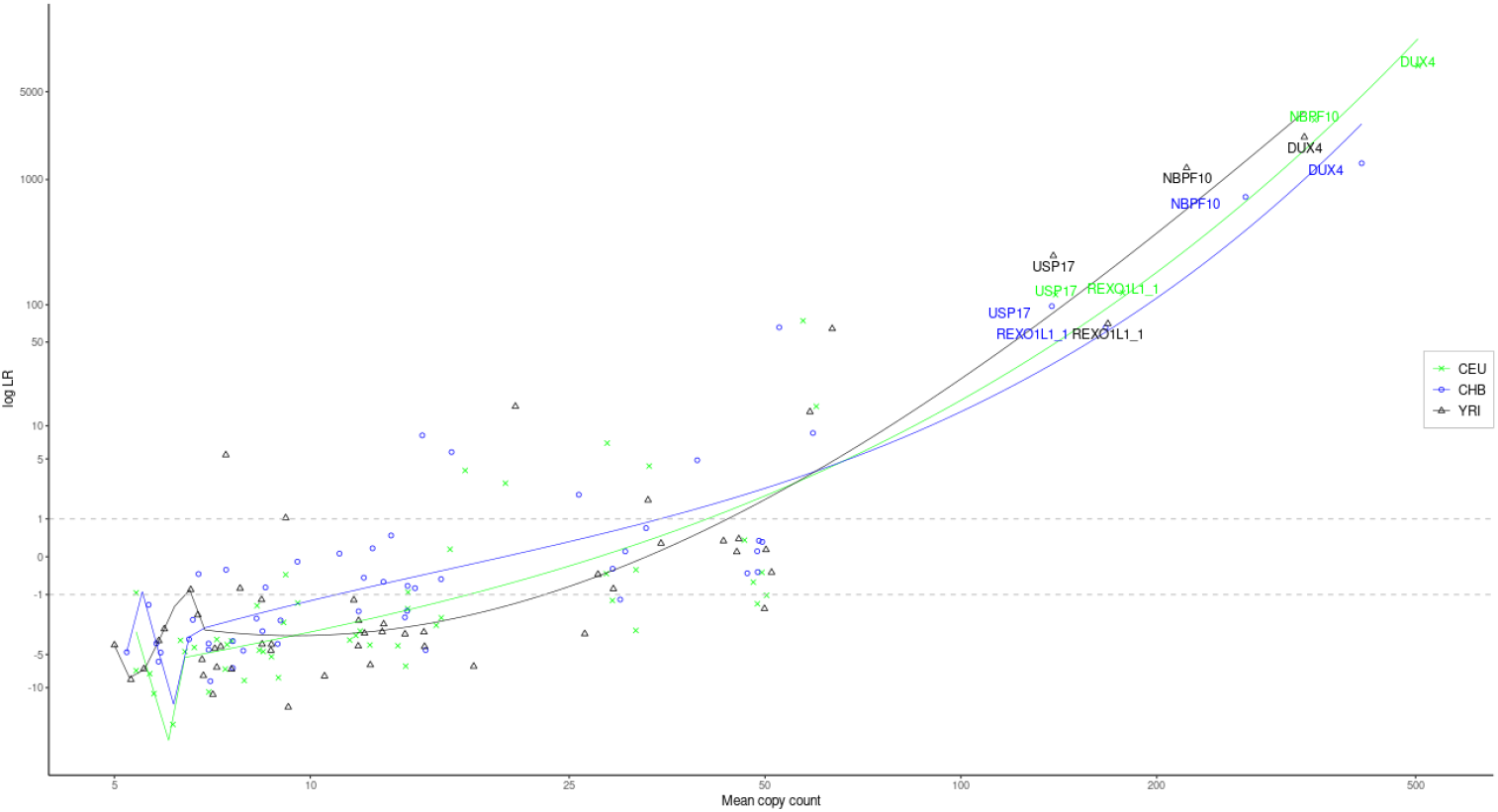
Comparison of average copy number with log likelihood ratios between CCM and STM fit of gene families observed in three human populations (YRI, CEU, CHB). We labelled all families with average copy numbers above 100. The fitted lines are spline interpolations (degree 3, knots at quantiles 0,0.05,0.1,…,1). The gray dotted lines highlight our cutoffs for positive evidence for either model (STM < −1, CCM > 1)

In general, there is a positive correlation between estimated *d/s* ratios and gene copy number both for gene families fitting better to STM and CCM across all populations (CCM: >0.89 (Pearson)/>0.56 (Spearman); STM: 0.71/>0.73 (Spearman)), even though not for all large gene families a high 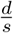 is estimated. In contrast, there is a clearly weaker correlation of core copy number (*i*_*o*_) estimates with mean copy numbers in CCM (Pearson: 0.016-0.54, Spearman: 0.35-0.73), compared to STM (Pearson: >0.95, Spearman: >0.77).

When we consider the variance to mean ratio, the maximum likelihood ratios do coincide well with the maximum likelihood estimates of *d/s*, i.e. the maximum likelihood differ at most by 24% from the variance to mean ratio (in CEU; ≤ 28%, ≤ 18% in CHB, YRI). This adds to the plausibility of CCM for these gene families. On the contrary, gene families with better fit to STM do not show a variance to mean ratio of approximately 1, but often much less (median/95%-quantile for variance to mean ration for CEU: 0.14/0.48, for CHB: 0.17/0.36, for YRI: 0.14/0.48). Thus, despite fitting the data better than CCM, it remains unclear if STM can truly explain the observed copy numbers in these cases.

## Discussion

We considered an evolutionary model where new copies of a gene arise by duplication and where selection favours chromosomes with low copy number. In the so-called single template model (STM) there is only one gene which can act as template for new copies. In the compound copy model (CCM) any copy may duplicate again. Re-interpreting mutations as new copies, STM is almost identical to Haigh’s (2) model of mutation-selection balance (when population size is infinite) and of Muller’s ratchet (when it is finite). We have shown that STM admits a finite equilibrium distribution with mean copy number *i*_*o*_ + *d/s*, where *i*_*o*_ is the number of ‘core’ copies – i.e., those which are present in all individuals – and *d/s* is the average number of accessible copies – i.e. those which are copy-number-variable in the population. Under the assumptions of infinite population size and absence of epistatic interactions this number is determined by the ratio of duplication rate *d* and strength of (purifying) selection *s*. For CCM we have shown that the limiting equilibrium distribution is essentially a negative binomial with parameters *i*_*o*_ and *s/*(*d* + *s*) and mean copy number *i*_*o*_(1 + *d/s*). The average number of accessory gene copies (the variable copy number added to the core copies) is given by the product *i*_*o*_ *d/s*.

For *i*_*o*_ = 1 both STM and CCM have on average the same number of extra copies. While remaining constant in STM, it linearly increases in *i*_*o*_ for CCM. For gene copy counts observed in a sample, one can assess the plausibility of the two models by simple means. From the equilibrium distributions of the two models follows that overdispersion (𝔽(***J*** ) > 1) is counterindicative of STM, while compatible with CCM. The parameters *d/s* and *i*_*o*_ can be estimated from moments of the hypothesized equilibrium distributions. One can also directly use a likelihood ratio approach for model selection and a maximum likelihood approach for parameter estimation. We illustrated this on two data sets of gene copy number variation (gCNV): (a) the NLR genes in *D. rerio* (19) and (b) several gene families in *H. sapiens* (17) (Figure 3, Tables 2,4). Counting zebrafish NLRs by their Fisnacht protein domains, and disregarding the lab population CGN, our estimates of the *d/s* ratio are roughly between 10 and 30 and the number of core copies (*i*_*o*_) between 6 (for TU) and 151 (for CGN). Both moment and maximum likelihood estimators agree very well in both estimates (Table 2). The CGN population, however, strongly differs from the other populations surveyed. Interestingly, the other lab population in our sample (TU) marks the opposite: *d/s* is estimated largest and *i*_*o*_ is estimated smallest among all populations. Again, very low effective population size, absence of random mating, and potentially a large copy number variance in the original founders could explain this observation. As noted before (19), lab populations are all but suitable to infer evolutionary properties of a species. Our results here confirm strong discrepancies in the immune gene repertoire between lab and wild populations of a model organism which is widely used in biomedical contexts to extrapolate research results to other vertebrates, including human.

As a second application, we analyzed in three populations of human a quite diverse set of gene families. They strongly differ in family size and range from very small to extremely large. Not surprisingly, copy number variation of small gene families is better fitted by STM, followed by a twilight zone where both models appear to be viable explanations of the observed gCNV, while large to super-large families can clearly be explained only by CCM. This mirrors our findings from large NLR gene copy numbers in *D. rerio*. In general, CCM appears to lead to better fits than STM. In small families, observed dispersion tends to be smaller than predicted under STM. Despite the good fit of CCM, in particular in large families, our model may be criticized for ignoring mechanisms which can play an important role in the evolution of gene copy numbers. Copy number can change by unequal recombination and duplicates can also arise by segmental or whole genome duplication. The latter are extremely rare events compared to those acting on a population genetic timescale of *O*(*N*_eff_) generations, and are neglected here. Our focus is on the copy numbers of individual gene families evolving within populations. As for the action of recombination we have previously analyzed a model where copy number change is induced by unequal cross-over (23) and shown that the equilibrium distribution is also very well approximated by a Gamma law. Therefore we do not expect fundamentally different predictions when recombination is included into model. However, to improve the accuracy of parameter estimation, it certainly remains desirable to integrate this mechanism. The drawback are the additional parameters which would have to be estimated or be determined by other means.

Another model assumption, which may be questioned, concerns the fitness function. The one considered here assumes a decrease in fitness with every additional gene copy. Insofar it is analogous to the fitness function of Haigh’s model, where any newly arising mutation is assumed to have a fitness cost. The question which fitness model is, or is not, appropriate has been discussed at length before (24–26). A one-fits-all answer does not exist. Rather, appropriateness depends not only on the size of the gene family but also on the genomic localization of the genes (arrayed in tandem or dispersed) and, above all, on gene function. The model from (26) treats selection on duplicates independently for each copy. This may be warranted for (very) small gene families, where different copies may have different functions, perhaps because they occupy different positions in a gene regulatory network or because they may be tuned to different environmental cues, for instance color vision genes in mammals (27, 28). Large gene families, for instance the NLRs considered as an example here, may comprise functionally active as well as pseudogenes. It certainly is conceivable that pseudogenes come with a fitness cost leading to decreasing overall fitness with increasing copy number. A unimodal fitness function may appear more appropriate when optimal fitness is associated with a certain fixed number of copies. However, similarly to Muller’s ratchet, low copy genotypes will eventually be lost under the action of drift, thereby transforming the fitness function from unimodal to monotonically decreasing. In any case, even if simplistic, the multiplicative fitness function, together with a tunable component of epistasis, covers a wide spectrum of ways how natural selection may shape gene copy number variation.

We explored by computer simulations the dynamics under finite population size, such that genetic drift induces a recurrent ratchet-like loss of the currently fittest class. In particular, we focused on the interarrival times between consecutive clicks, or the speed, of the ratchet. In CCM duplication is a self-accelerating process leading to ever decreasing, and ultimately vanishing, interarrival times. Such a process may be called run-away evolution. With very different model assumptions the generation of large families by fast spread of copies in a population has also been the subject of a recent study by (29).

By a very natural generalization of the fitness function, our models provide for including the effect of epistasis on the evolutionary dynamics of multi-copy gene families. One of the most striking properties of synergistic epistasis is its capacity to slow down the speed of the ratchet. Synergistic epistasis (*β* > 1) induces strong negative selection against individuals with high copy numbers and acts like a barrier against additional copies. Only once surmounted, the ratchet moves with accelerating speed and without upper bound on average copy number. Thus, depending on the strength of epistasis *β* and the relative magnitudes of *d* and *s*, interarrival times may show a transient pattern: increasing while accessory copy number *i* is small, but then decreasing and eventually vanishing when *i* is sufficiently large (Figure 2). Under an extreme form of synergistic epistasis – truncation selection – where individuals with more than a (finite) threshold number of copies have fitness 0, the ratchet ceases to click when all individuals have reached the threshold.

An ever increasing copy number does not seem plausible in practice. Even in large gene families, such as the NLRs in zebrafish, gene copy number variation appears to be smaller than expected under a regime of pure run-away evolution. Our simulations show that epistasis can substantially slow down the ratchet already in very small (here *N*_eff_ = 400) to moderately large (here *N*_eff_ = 2000) populations while other parameters are kept in a plausible range (e.g., *s* = 2*e*-4, *d* = 2*e*-3, corresponding to *d/s* = 10, see Fig 2). In both cases we observe an increase of the interarrival times between ratchet clicks in the interval roughly between *k* = 100 and *k* = 1000. Using different arguments, and extrapolating from observed copy counts, Schäfer et al. (19) estimated upper bounds on the total number of NLR genes in zebrafish populations and arrived at values of a few thousand for wild populations and of a few hundred for the lab strains (see their Table 1 in (19)). Thus, even if natural populations may not (yet) be at a stable equilibrium, they may have entered a metastable state with interarrival times way beyond a population genetic time scale of *O*(*N*_eff_) generations (Fig 2).

At large, the same arguments can be extended to interpreting copy number variation of large gene families in human populations in the light of our model. However, as noted before, CCM may not apply to small gene families. If variable at all, their dynamics may well be explained by STM. Here, run-away evolution is not an issue.

## ACKNOWLEDGEMENTS

This work was supported by (separate) grants to TW and FF from the German Research Society (DFG) in the framework of SPP-1590 and SPP-1819.

## Data availability

Simulation data are available at https://github.com/y-zheng/Runaway-Duplication-Simulations/. R- code used for numerical analyses is available at https://github.com/fabfreund/genedup. Experimental data from *D. rerio* and human are available in the cited references.

## Supplementary Material

### Suppl 1. Convergence to equilibria in the infinite case

In this section, we first present all results and the general arguments we use to establish convergence of both mean fitness and of the copy count frequencies in STM and CCM under infinite population size and no epistasis (*β* = 1). All proofs then follow in the subsequent section. To shorten the calculations, we define

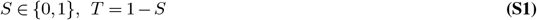

where *S* is an indicator variable of whether CCM is assumed; that means, under CCM we have *S* = 1 whereas *S* = 0, and hence *T* = 1, under STM.

We will first show convergence of the mean fitness (Proposition 2). The limit does not depend on the strength of selection.

#### Proposition 2.

*In the case of infinite population size, the mean fitness converges to*

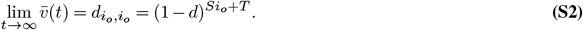

In the following, convergence of the relative frequencies (*y*_0_(*t*), *y*_2_(*t*),…) in the vector space 𝓁^1^ will be established. Let 𝓁 denote the field of all vectors (*z*_0_, *z*_1_, …) ∈ 𝓁^1^ with *z*_0_ = 1. Let *y*(*t*) = (*y*_0_(*t*), *y*_1_(*t*),…) denote a vector of relative frequencies of copy numbers under STM or CCM. Then

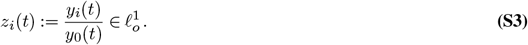

Both STM and CCM satisfy *y*_0_(*t*) > 0 ∀*t* in the infinite case. Considering the time dynamics of *z*_*i*_(*t*), we have

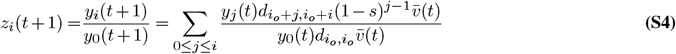

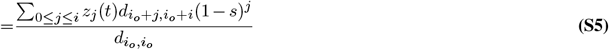

Since Σ_*i*≥0_ *y*_*i*_ = 1, we have ∥*z*(*t*)∥_1_ := Σ_*i*≥0_ *z*_*i*_(*t*) = (*y*_0_(*t*))^−1^ > 0. Also, any 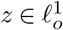 can be transformed into a probability distribution on ℕ_0_ by setting 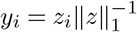 for *i* ≥ 0.

#### Lemma 1.

*For any starting frequencies* (*y*_*i*_(0))_*i*≥0_ *in STM and CCM, the corresponding* 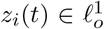 , *t ∈ ℕ*_0_, *converge component-wise:*

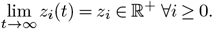

In order to prove this, we will need a simple result on convergence of recursively defined sequences:

#### Lemma 2.

*Let* (*γ*_*i*_)_*i∈*ℕ_ *be a convergent sequence of real numbers with limit γ* ∈ ℝ. *Let x*_0_ ∈ ℝ, *a* ∈ (0, 1) *and the series* (*x*_*i*_)_*i∈*ℕ_*be recursively defined by*

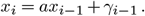

*Then*

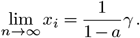

To show convergence in 𝓁^1^ of the transformed system (*z*(*t*))_*t*_≥_0_ and subsequently also of the non-transformed system (*y*(*t*))_*t*_, we need to control the tail of the distribution of *y*(*t*) for *t* large.

#### Lemma 3.

*For any system* (*y*(*t*))_*t*_ *in either STM or CCM, the tail probabilities are vanishing for large t, i*.*e. there exist t*_0_, *k*_0_ *∈* ℕ *so that*

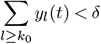

*for any δ* > 0 *and t* ≥ *t*_*o*_.

With that, we can show

#### Lemma 4.

*Consider an arbitrary starting configuration y*(0) *in STM or CCM and let* 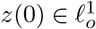 *be the corresponding transformed vector. Then*, 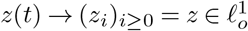 *for t* → ∞, *where for i ∈* ℕ_0_, *z*_*i*_ *is the limit defined in Lemma 1. z is the unique fixed point of* 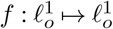 *defined by*

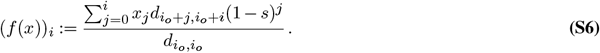

From this, an analogous result follows directly for the non-transformed system *y*(*t*), proving Proposition 1.

### Suppl 2. Propositions and Lemmata

*Proof of Proposition 2*. To simplify notation, we introduce the (generating) function

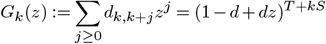

for the binomial duplication probabilities in STM and CCM respectively. Consider additionally the following recursive set of expressions:

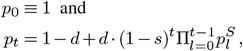

where *S* is defined as in Eq. (S1). We may interpret these also as polynomials *p*_*t*_(*s*) of the selection coefficient. We have *p*_*t*_ ∈ (0, 1] for *t* ∈ ℕ_0_, and thus *p*_*t*_ → 1−*d* for *t*→∞. Considering Eq. (6), the mean fitness *after* duplication, selection and normalisation amounts to

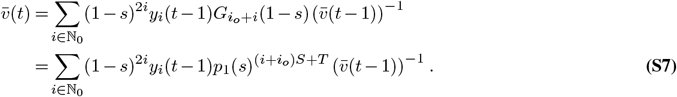

Expressing *y*_*i*_(*t*− 1) in terms of *y*_*j*_(*t*− 2), *j* ≤ *i* and applying Eq. (6), we obtain:

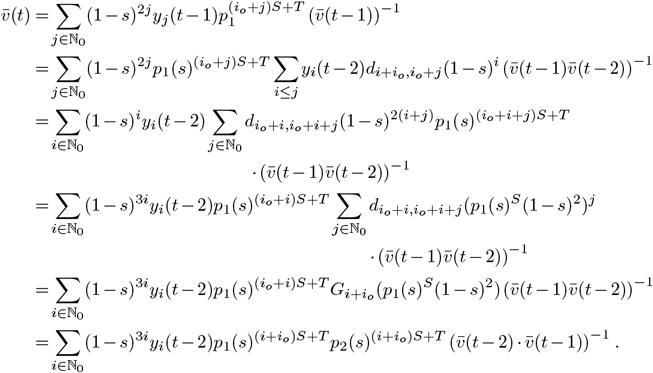

Repeating this *t*− 2 times, one arrives at

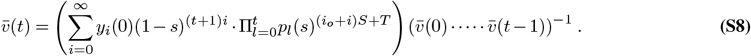

Using the same procedure to rewrite the denominator, one finally obtains

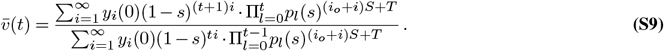

Consider first the starting configuration given by the vector *y* = *y*^(^0) = (1, 0,… ) ∈ 𝓁_1_. Since *y*_*i*_(0) = 0 for all *i* > 1, Eq. (S9) reads:

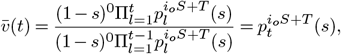

which converges to 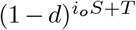 for *t* → ∞.

Now, consider a slightly more general initial configuration *y*^(0)^ *∈* 𝓁_1_ with 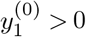. By Eq. (S9), we may write

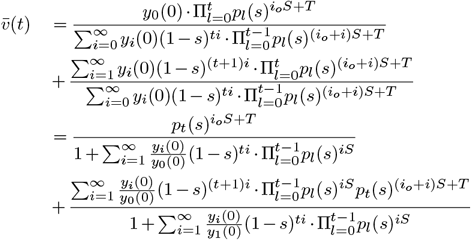

Since *y*(0) *∈* 𝓁_1_, the sum 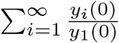 converges. Because |*p*_*l*_(*s*)| < 1 for all *l*, and because (1 −*s*)^*ti*^ and (1 −*s*)^(*t*+1)*i*^ converge to 0, we have

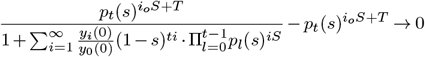

and

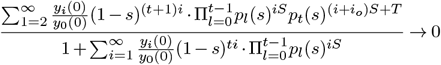

This shows that the mean fitness 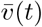 converges to 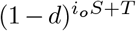 for any initial *y* ^(0)^ with 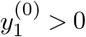.

*Proof of Lemma 2*. Since 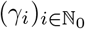 is convergent, it is also bounded, i.e. there exists a constant *B* > 0 such that |*γ*_*i*_| *< B* for all *n ∈* ℕ. For any given *ϵ* > 0, we may choose *N ∈* ℕ large enough such that

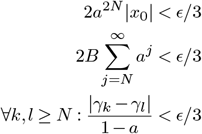

The last statement is true because 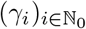 is a Cauchy sequence.

Iterating the recursive formula for *x*_*i*_ shows that 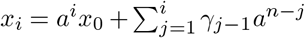. Now let *n > m* be any two natural numbers, then the following holds:

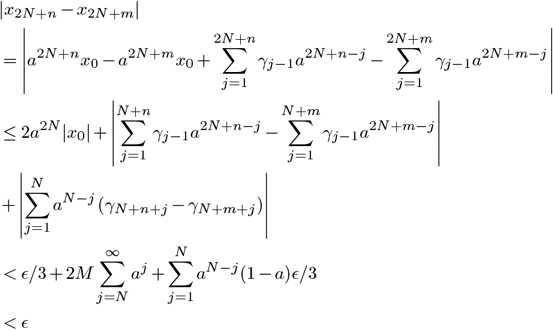

Thus, 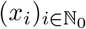 is also a Cauchy sequence and therefore convergent. Since the recurrence relation must also hold for the limit, i.e.

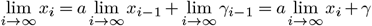

we obtain, after rearranging,

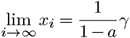

*Proof of Lemma 1*. By construction, *z*_0_(*t*) ≡ 1, which trivially converges for *t* → ∞. Eq. (S5) shows that for *i* > 0, *z*_*i*_(*t* + 1) is a linear combination, say *γ*_*t*+1_, of finitely many *z*_*j*_(*t*), 0 ≤ *j < i* with positive coefficients which do not depend on *t*, and of *z*_*i*_(*t*) with coefficient (1 − *d*)^*iS*^ (1 − *s*)^*i*^. Suppose the sequences (*z*_*j*_(*t*))_*t∈*ℕ_ are convergent for *j ∈* {0,…, *i*− 1} to positive limits. Then, also the linear combinationy *γ*_*t*_ converge to a positive limit *γ* for *t* → ∞. Thus, (*z*_*i*_(*t*))_*t∈*ℕ_ is convergent after application of Lemma 2 with *a* = (1 − *d*)^*iS*^ (1 − *s*)^*i*^ < 1 and *γ*_*i*_ defined as described above. Moreover, since *γ* > 0, Lemma 2 ensures lim_*t*→∞_ *z*_*i*_(*t*) = (1 −*a*)^−1^*γ* > 0. The lemma now follows from induction.

*Proof of Lemma 3*. Recall the asymptotic mean fitness 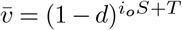. As thefitnesses *v*(*t*) at times *t ∈* ℕ_0_ converge, we can choose *t*_0_ so that 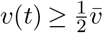 for *t* ≥ *t*_0_. Consider now the tail probability *τ*_*k*_ = Σ_*l*≥*k*_ *y*_*l*_(*t* + 1) for *t* ≥ *t*_0_.

Observe that only mass from 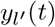 for 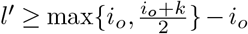 gets shifted to *y*_*l*_(*t* + 1), *l* = *k, k* + 1, *k* + 2,… at time *t* + 1 by duplication and thus contributes to the tail probability of *y*(*t* + 1). Due to selection, any *y*_*l*_(*t*) gets weighted by 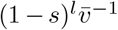. This means for *k* bigger than *i*_*o*_, let’s say *k* = *k*^*′*^ + *i*_*o*_ with *k*^*′*^ > 0, any mass relevant for the tail probability has a weight bounded by 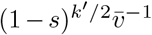. Since the mass in *y*(*t*) equals 1, this means that 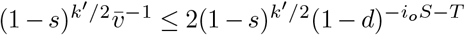, using the fitness bound established above. This gets arbitrarily small for *k*^*′*^ large enough.

*Proof of Lemma 4*. Since ∥*y*(*k*)∥_1_ = 1 for all *t*, Lemma 3 is equivalent to 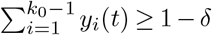 for *t > k* for adequately chosen *L*. Thus, we have

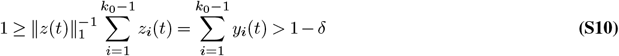

for *t > K*. Lemma 1 shows that 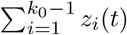 converges to a positive limit, so Eq. (S10) implies that (∥*z*(*t*)∥_1_)^*t∈*ℕ^ has finite and positive upper and lower bounds. Moreover, this boundedness shows that for any *ϵ* > 0 we can find *L*^*′*^and *t*^*′*^so that 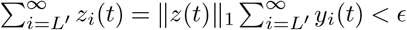 for all *t* ≥ *t*^*′*^, which implies convergence of ∥*z*(*t*)∥ for *t*→ ∞. This ensures also 𝓁^1^ convergence of (*z*(*t*))_*k∈*ℕ_ to the limit *z* = (*z*_*i*_)_*i∈*ℕ_, to which pointwise convergence was established in Lemma 1. This follows since for sufficiently large *N* and *k* → ∞, all three sums on the right hand side of the inequality

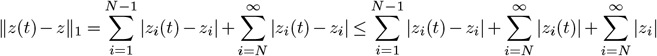

become arbitrarily small.

It remains to be shown that this limit is unique. Eq. (S5) shows that *z*(*t* + 1) = *f* (*z*(*t*)). If *f* is continuous, we have

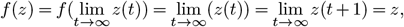

and the uniqueness follows from the convergence to *z*. Thus, we only need to show that *f* is continuous. We do this by showing Lipschitz continuity. For *x, x*^*′*^ *∈ 𝒳*_*o*_ we have

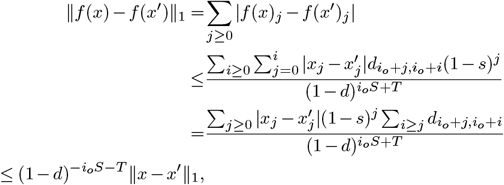

where we use the fact that the duplication probabilities in a state with *j* copies sum up to 1.

*Proof of Proposition 1*. Since 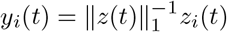, Lemma 4 implies 𝓁^1^-convergence of *y*(*t*) for *t* → ∞, from which the reverse triangle inequality also implies that ∥*y*^*^∥_1_ = 1. Since ∥*z*(*t*)∥_1_ converges for *t* → ∞, the fixed point equation for (*z*(*t*))_*t∈*ℕ_ remains valid as is for (*y*(*t*))_*t∈*ℕ_, i.e. *f* (*y*^*^) = *y*^*^. Eq. (11) follows from Eq. (S6) solving for the *j*th (resp. *i*th) coordinate of the fixed point.

### Suppl 3. Approximation quality of equilibria distributions

While we cannot directly compare the recursive description of the equilibrium frequencies in Eq. (8) with the Poisson and negative binomial approximations, we can compare the conditional distributions/frequency of individuals with *I* (extra) gene copies among individuals with at most *N* copies, i.e.

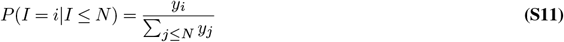

for 0 ≤ *i* ≤ *N* and = 0 for *i > N* to the associated conditional Poisson and negative binomial approximations.

It is straightforward to compute these conditional probabilities both for the exact equilibria (Eq. (S11), via the recursion Eq. (8)) and for the approximations (our R implementation can be found here: tba). Moreover, we can also compute *P* (*I*≤*N* ) for the approximations and a lower bound for it for the exact equilibrium. For the latter, we simply observe, with the mean fitness at equilibrium being 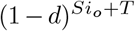,

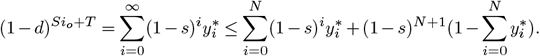

Note that we can express the right hand side as a function of 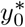 due to the recursive definition of thee 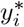’s. We denote the relation between 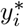 and 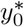 by 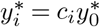 Then, if we treat this inequality as an equality, we can solve for an unique solution 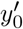 that satisfies 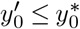. Then, we have the lower bound 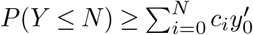.

See Table S1 for a comparison between the exact conditional distribution and their approximations.

### Suppl 4. Bonferroni-type correction for likelihood ratios

Consider comparing two hypotheses *H*_1_, *H*_2_ by their likelihoods of observed data *D*. In a Bayesian framework with equal prior probabilities of 0.5, this means that the odds ratio (OR) equals the likelihood ratio (LR), i.e.

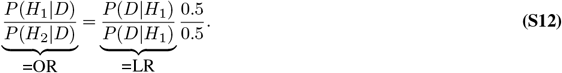

Without restriction, assume that LR>1, so *H*_2_ is the less likely hypothesis in the Bayesian approach. With two hypotheses, we have *P* (*H*_1_ |*D*) + *P* (*H*_2_ |*D*) = 1. Thus, we can use Eq. (S12) to compute the posterior probability of the less likely hypothesis *H*_2_ as

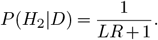

As a test criterion, we can control this posterior probability, similar to a frequentist significance level, which is the thought behind the Kass-Raftery scale (30). Indeed, allowing at most a significance level of *α*, leads to the condition *LR* ≥ α^−1^ −1, and thus log *LR* ≥ log(α^−1^ −1), which for *α* = 0.05 equals approx. 2.944, which then is typically rounded up to 3, for a slightly conservative threshold (which is called ßtrong evidenceïn the Kass-Raftery scale). In case of LR<1, we switch hypotheses, which leads to the same threshold with a negative sign (for *α* = 0.05, this explains the treshold of −3 for strong evidence).

We can thus easily expand that to control the overall posterior probability for the ‘other’ hypothesis across all comparisons by simply applying a Bonferroni correction to *α*, so computing the thresholds with *α*^*′*^ = *α/n* where *n* is the number of comparisons performed.

## Supplementary Tables

**Table S1.** Conditional distributions of (extra) copy numbers *I* at equilibria frequencies, conditioned on *I* ≤ 800, for various parameter contributions and their approximations by appropriate Poisson and negative binomial distributions (with same conditioning). exc: exact conditional distribution, appx: conditional approximated distribution, t.v.d: total variation distance, ||.||_∞_: maximal distance between exc and appx for given parameters. See Sect. Suppl 3 for details. We have *P* (*I* ≤ 800) = 1 for the conditional exact distribution and its approximation for all parameter choices.

| Parameters | mean exc | mean appx | s.d. exc | s.d. appx | $\vert \vert \cdot\vert \vert _\infty$ | t.v.d. |
| --- | --- | --- | --- | --- | --- | --- |
| CCM $i_o = 1, d = 0.01, s = 10^{-3}$ | 10.100 | 10.0 | 10.536 | 10.488 | 0.002 | 0.005 |
| CCM $i_o = 1, d = 10^{-3}, s = 10^{-4}$ | 10.010 | 10.0 | 10.493 | 10.488 | 0.000 | 0.000 |
| CCM $i_o = 25, d = 0.01, s = 10^{-3}$ | 252.511 | 250.0 | 52.679 | 52.440 | 0.000 | 0.019 |
| CCM $i_o = 25, d = 10^{-3}, s = 10^{-4}$ | 250.250 | 250.0 | 52.464 | 52.440 | 0.000 | 0.002 |
| CCM $i_o = 1, d = s = 10^{-3}$ | 1.001 | 1.0 | 1.415 | 1.414 | 0.000 | 0.000 |
| CCM $i_o = 1, d = s = 10^{-4}$ | 1.000 | 1.0 | 1.414 | 1.414 | 0.000 | 0.000 |
| CCM $i_o = 25, d = s = 10^{-3}$ | 25.025 | 25.0 | 7.073 | 7.071 | 0.000 | 0.001 |
| CCM $i_o = 25, d = s = 10^{-4}$ | 25.003 | 25.0 | 7.071 | 7.071 | 0.000 | 0.000 |
| CCM $i_o = 1, d = 10^{-4}, s = 10^{-3}$ | 0.100 | 0.1 | 0.332 | 0.332 | 0.000 | 0.000 |
| CCM $i_o = 1, d = 10^{-5}, s = 10^{-4}$ | 0.100 | 0.1 | 0.332 | 0.332 | 0.000 | 0.000 |
| CCM $i_o = 25, d = 10^{-4}, s = 10^{-3}$ | 2.500 | 2.5 | 1.658 | 1.658 | 0.000 | 0.000 |
| CCM $i_o = 25, d = 10^{-3}, s = 10^{-4}$ | 2.500 | 2.5 | 1.658 | 1.658 | 0.000 | 0.000 |
| STM $i_o = 1, d = 0.01, s = 10^{-3}$ | 10.050 | 10.0 | 3.162 | 3.162 | 0.001 | 0.007 |
| STM $i_o = 1, d = 10^{-3}, s = 10^{-4}$ | 10.005 | 10.0 | 3.162 | 3.162 | 0.000 | 0.001 |
| STM $i_o = 25, d = 0.01, s = 10^{-3}$ | 10.050 | 10.0 | 3.162 | 3.162 | 0.001 | 0.007 |
| STM $i_o = 25, d = 10^{-3}, s = 10^{-4}$ | 10.005 | 10.0 | 3.162 | 3.162 | 0.000 | 0.001 |
| STM $i_o = 1, d = s = 10^{-3}$ | 1.001 | 1.0 | 1.000 | 1.000 | 0.000 | 0.000 |
| STM $i_o = 1, d = s = 10^{-4}$ | 1.000 | 1.0 | 1.000 | 1.000 | 0.000 | 0.000 |
| STM $i_o = 25, d = s = 10^{-3}$ | 1.001 | 1.0 | 1.000 | 1.000 | 0.000 | 0.000 |
| STM $i_o = 25, d = s = 10^{-4}$ | 1.000 | 1.0 | 1.000 | 1.000 | 0.000 | 0.000 |
| STM $i_o = 1, d = 10^{-4}, s = 10^{-3}$ | 0.100 | 0.1 | 0.316 | 0.316 | 0.000 | 0.000 |
| STM $i_o = 1, d = 10^{-5}, s = 10^{-4}$ | 0.100 | 0.1 | 0.316 | 0.316 | 0.000 | 0.000 |
| STM $i_o = 25, d = 10^{-4}, s = 10^{-3}$ | 0.100 | 0.1 | 0.316 | 0.316 | 0.000 | 0.000 |
| STM $i_o = 25, d = 10^{-3}, s = 10^{-4}$ | 0.100 | 0.1 | 0.316 | 0.316 | 0.000 | 0.000 |

**Table S2.** Sequence read numbers and exon counts of FISNA-NACHT and NLR-associated B30.2 exons from experimental data.

| **Population**<br>**(diploid**<br>**sample size)** | **CGN (8)** | **CHT (18)** | **DP (20)** | **KG (20)** | **SN (19)** | **TU (8)** |
| --- | --- | --- | --- | --- | --- | --- |
| **Reads** | 18935,12851,<br>11853,17235,<br>26914,26200,<br>23001,24546 | 78648,142018,<br>102969,117377,<br>75181,117936,<br>80986,90855,<br>85293,87768,<br>79771,9953,<br>52384,144266,<br>39048,49949,<br>69193,57095 | 790,8166,<br>44471,26848,<br>25214,29932,<br>9091,8683,<br>10686,143954,<br>12269,8313,<br>7376,8658,<br>20187,9913,<br>7555,13210,<br>7764,4612 | 43957,42564,<br>60852,36727,<br>32824,32297,<br>44678,27109,<br>37385,45094,<br>50191,39644,<br>42213,33860,<br>57148,37160,<br>18606,17986,<br>26969,32098 | 129537,63760,<br>43631,55884,<br>111227,55034,<br>30486,66059,<br>94355,114889,<br>106075,75794,<br>108626,92966,<br>122603,51335,<br>92094,222595,<br>59689 | 18886,17606,<br>24125,26990,<br>12495,9358,<br>5338,8028 |
| **FISNACHT**<br>**exons** | 269,267,299,<br>264,271,245,<br>284,276 | 387,532,418,<br>459,418,410,<br>389,410,420,<br>404,381,167,<br>348,534,305,<br>355,374,383 | 61,278,336,<br>285,286,294,<br>258,250,298,<br>435,279,282,<br>261,252,328,<br>289,260,309,<br>235,201 | 485,444,518,<br>382,402,408,<br>477,351,431,<br>429,448,445,<br>464,454,496,<br>282,231,223,<br>280,365 | 521,399,385,<br>405,456,396,<br>344,534,468,<br>562,543,399,<br>443,450,491,<br>335,378,524,<br>358 | 238,229,246,<br>264,120,126,<br>78,112 |
| **NLR-B30.2**<br>**exons** | 88,79,77,<br>73,90,85,<br>88,88 | 128,174,137,<br>164,143,143,<br>134,139,140,<br>147,122,66,<br>121,174,86,<br>101,117,102 | 12,57,75,<br>67,60,60,<br>71,51,65,<br>134,61,61,<br>55,63,79,<br>61,48,76,<br>55,46 | 159,161,180,<br>137,143,137,<br>162,118,141,<br>147,164,154,<br>161,160,180,<br>75,60,61,<br>73,129 | 153,126,143,<br>120,142,121,<br>108,169,146,<br>187,160,129,<br>118,128,144,<br>102,113,153,<br>109 | 68,73,76,<br>73,35,45,<br>28,35 |

**Table S3.** Human gene families with minimum observed copy number in three populations, YRI, CEU and CHB. Original data from (17), adapted by (23). Gene families with zero variance (no copy number variation) in either of the populations were excluded from analysis.

| ID | Name | YRI | CEU | CHB | ID | Name | YRI | CEU | CHB | ID | Name | YRI | CEU | CHB |
| --- | --- | --- | --- | --- | --- | --- | --- | --- | --- | --- | --- | --- | --- | --- |
| 1 | AMY1A | 5 | 4 | 5 | 52 | GOLGA8G | 22 | 26 | 27 | 103 | PLGLB2 | 7 | 7 | 7 |
| 3 | ANKRD20A3 | 21 | 25 | 26 | 53 | GSTM | 2 | 2 | 2 | 104 | PPIAP21 | 35 | 38 | 43 |
| 4 | ANXA8L1 | 3 | 3 | 4 | 54 | GSTT2 | 2 | 2 | 2 | 105 | PRAMEF14 | 7 | 7 | 8 |
| 6 | ARL17B | 3 | 3 | 3 | 55 | GTF2H2C | 3 | 3 | 3 | 106 | PRAMEF20 | 6 | 6 | 5 |
| 8 | AURKAIP1 | 2 | 2 | 2 | 56 | GUSBP1 | 8 | 10 | 9 | 107 | PRAMEF5 | 12 | 12 | 11 |
| 9 | BOLA2B | 5 | 5 | 5 | 58 | HIST2 | 7 | 7 | 7 | 108 | PRAMEF8 | 5 | 3 | 4 |
| 10 | BTNL8 | 2 | 2 | 2 | 59 | HP | 2 | 2 | 2 | 109 | PRR11 | 5 | 7 | 7 |
| 11 | C16orf54 | 2 | 2 | 2 | 60 | HRNR | 1 | 1 | 1 | 110 | PRR20A_2 | 8 | 7 | 7 |
| 12 | C4B | 2 | 3 | 3 | 61 | KIF22 | 2 | 2 | 2 | 111 | PRR20A_1 | 8 | 6 | 7 |
| 14 | CBWD3 | 10 | 10 | 10 | 62 | KIR2DL2 | 7 | 7 | 8 | 112 | PSG3 | 12 | 13 | 10 |
| 15 | CCL3 | 2 | 0 | 1 | 63 | KIR2DL4 | 0 | 3 | 2 | 113 | PTPN20A | 3 | 3 | 3 |
| 16 | CCL4 | 3 | 2 | 3 | 64 | KIR2DL5A | 7 | 6 | 7 | 114 | PYCR1 | 2 | 2 | 1 |
| 17 | CCR2 | 2 | 2 | 2 | 65 | KIR2DS2 | 6 | 7 | 7 | 115 | REXO1L1_1 | 126 | 110 | 117 |
| 20 | CDC37P1 | 5 | 9 | 6 | 66 | KIR3DP1 | 0 | 6 | 4 | 116 | REXO1L1_2 | 108 | 128 | 119 |
| 22 | CKMT1A | 3 | 3 | 4 | 67 | KLRC2 | 3 | 3 | 3 | 117 | RGPD1 | 12 | 10 | 12 |
| 23 | CLEC18A | 6 | 6 | 5 | 68 | KPNA2 | 3 | 3 | 3 | 118 | RGPD6 | 13 | 12 | 13 |
| 24 | CR1 | 3 | 3 | 3 | 71 | KRTAP1-1 | 1 | 1 | 1 | 119 | RPS17 | 2 | 2 | 2 |
| 25 | CSH | 6 | 6 | 7 | 73 | KRTAP3-1 | 2 | 3 | 4 | 120 | RSPH10B | 3 | 3 | 4 |
| 26 | D2HGDH | 2 | 2 | 2 | 74 | KRTAP4-1 | 1 | 1 | 2 | 121 | SCRT1 | 1 | 1 | 2 |
| 27 | DEFA1 | 4 | 4 | 4 | 78 | KRTAP9-1 | 3 | 3 | 3 | 122 | SCXB | 0 | 1 | 0 |
| 28 | DEFB103A | 2 | 2 | 2 | 79 | LCE2 | 3 | 3 | 3 | 123 | SERF1B | 2 | 2 | 2 |
| 29 | DEFB104A | 3 | 2 | 2 | 80 | LIMS3 | 5 | 5 | 5 | 124 | SFTPA1 | 3 | 3 | 3 |
| 30 | DEFB105A | 1 | 2 | 2 | 81 | LOC100506776 | 2 | 1 | 2 | 125 | SIRT7 | 2 | 2 | 2 |
| 31 | DEFB106A | 2 | 2 | 2 | 82 | LOC23117 | 43 | 42 | 44 | 126 | SLX1B | 5 | 4 | 4 |
| 32 | DEFB107A | 2 | 2 | 2 | 83 | LOC653606 | 6 | 5 | 4 | 127 | SMN2 | 2 | 2 | 2 |
| 33 | DEFB130 | 4 | 4 | 4 | 84 | LOC727800 | 1 | 1 | 1 | 128 | SNAR-A1 | 32 | 37 | 32 |
| 34 | DGAT1 | 2 | 2 | 2 | 85 | LOC728932 | 3 | 3 | 4 | 129 | SPAG11 | 2 | 2 | 2 |
| 35 | DHRS4 | 4 | 4 | 4 | 86 | LPA | 2 | 4 | 2 | 130 | SPDYE3 | 29 | 25 | 29 |
| 37 | DUX4 | 200 | 212 | 259 | 87 | MAFG | 2 | 2 | 2 | 131 | SULT1A1 | 3 | 2 | 2 |
| 38 | EIF3C | 3 | 3 | 3 | 88 | MGC59937 | 2 | 2 | 2 | 132 | SULT1A3 | 5 | 5 | 5 |
| 39 | ELA | 3 | 3 | 3 | 90 | MUC12 | 6 | 10 | 9 | 133 | TAS1R3 | 2 | 2 | 2 |
| 40 | FAM203A | 2 | 2 | 2 | 91 | MUC5AC | 2 | 2 | 2 | 134 | TBC1D3_1 | 32 | 21 | 24 |
| 41 | FAM21B | 4 | 4 | 4 | 92 | NBPF10 | 126 | 154 | 179 | 135 | TBC1D3_2 | 32 | 20 | 23 |
| 43 | FAM72A | 5 | 6 | 6 | 93 | NBPF11 | 38 | 40 | 41 | 136 | TCEB3C | 16 | 11 | 12 |
| 44 | FAM75A1 | 9 | 9 | 10 | 94 | NBPF16 | 36 | 37 | 37 | 140 | TMPRSS11E | 1 | 1 | 2 |
| 45 | FAM75A5 | 10 | 9 | 10 | 95 | NDUFB4 | 2 | 2 | 2 | 141 | TP53TG3 | 4 | 4 | 4 |
| 46 | FAM90A | 24 | 19 | 21 | 96 | NEB | 1 | 1 | 2 | 142 | TRIM49L1 | 8 | 10 | 10 |
| 47 | FCGBP | 3 | 4 | 4 | 98 | NPIP | 47 | 44 | 45 | 143 | UGT2B15 | 2 | 1 | 2 |
| 48 | FCGR | 2 | 3 | 2 | 99 | ORM1 | 3 | 4 | 3 | 144 | USP17 | 74 | 83 | 77 |
| 49 | FOXD4L2 | 11 | 11 | 11 | 100 | P4HB | 2 | 2 | 2 | 145 | ZFP62 | 1 | 1 | 1 |
| 50 | FUSIP1 | 2 | 2 | 2 | 101 | PCYT2 | 2 | 2 | 2 | 146 | ZG16 | 1 | 2 | 1 |
| 51 | GOLGA6L9 | 23 | 23 | 24 | 102 | PGA3 | 4 | 2 | 6 | 147 | ZNF658B | 4 | 5 | 5 |

## Supplementary Figures

**Figure S1.**
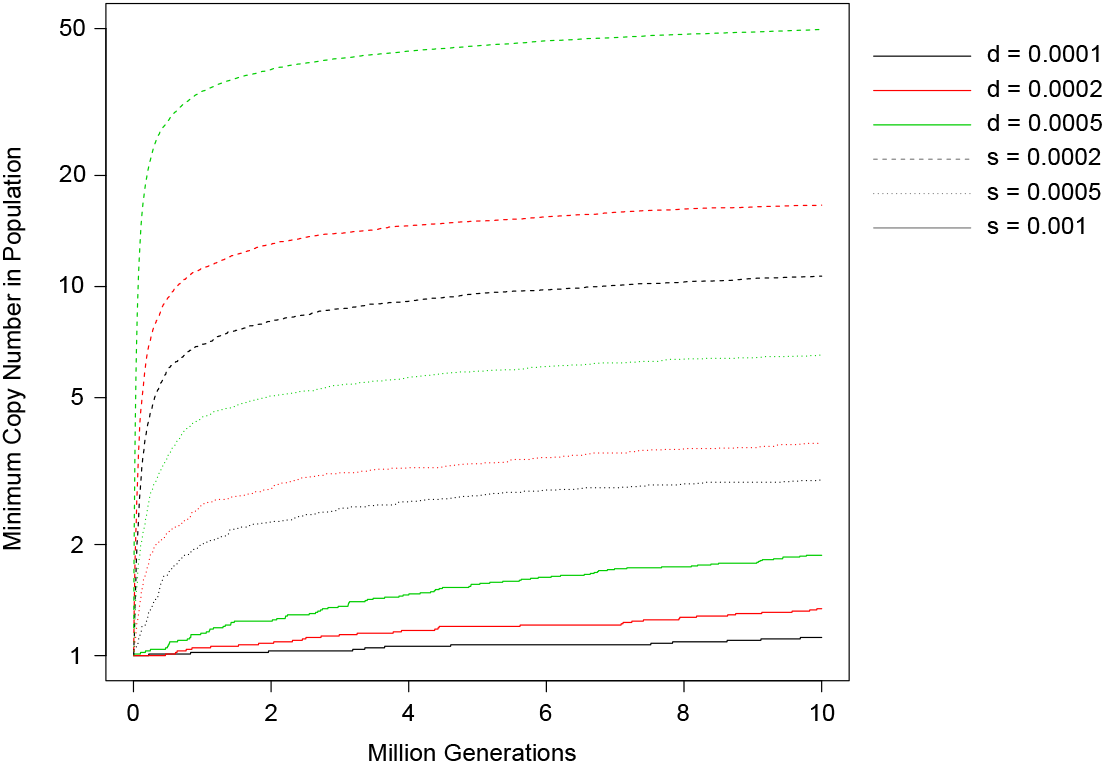
Core copy number *vs*. time. Average across 100 replicates. CCM with *β* = 2. For each line, *d* is coded by color and *s* coded by line type (see legend).

**Figure S2.**
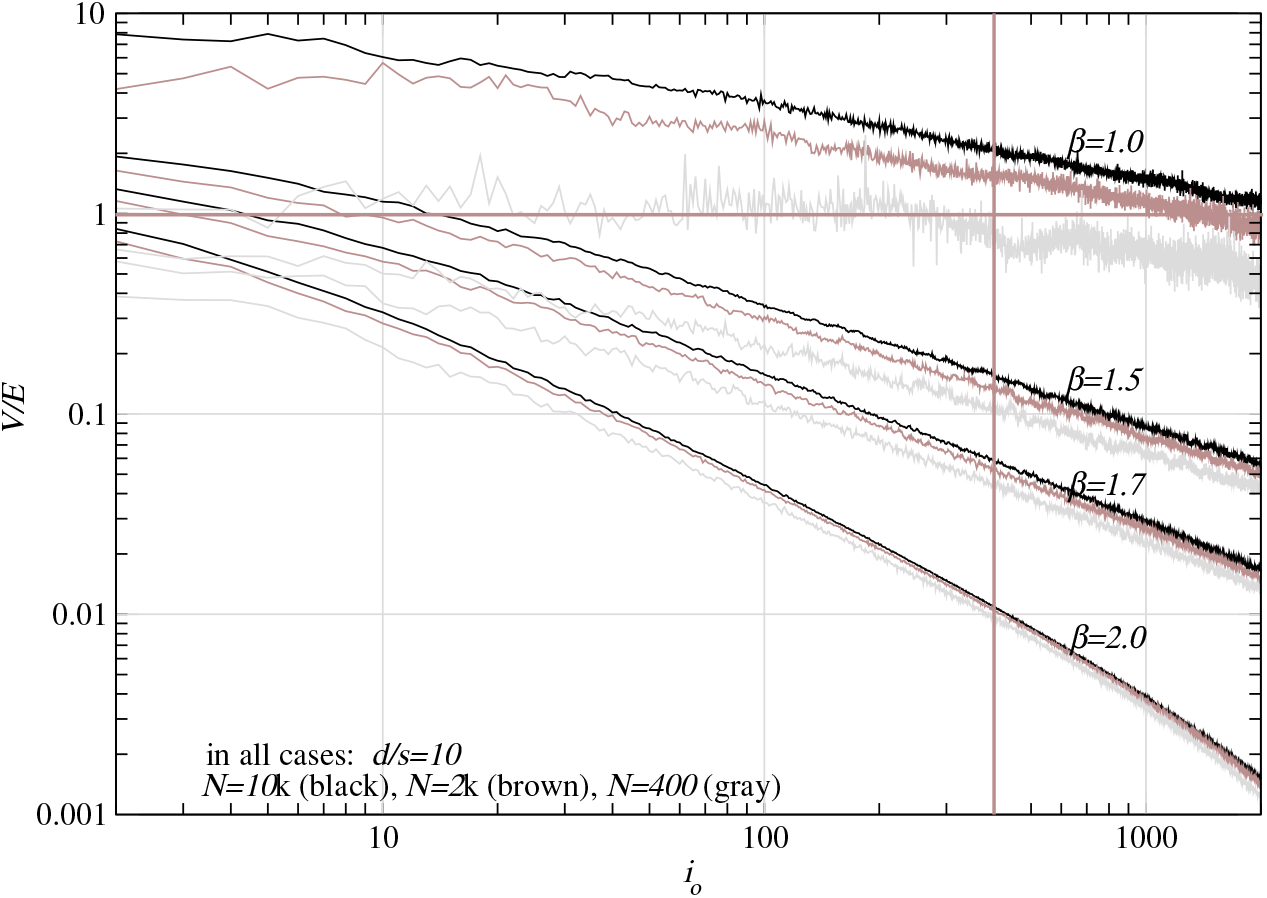
Index of dispersion (𝔽 = 𝕍*/*𝔼) in copy number for CCM model without (*β* = 1.0) and with (*β* = 1.5, 1.7, 2.0) epistasis. Epistasis reduces dispersion. *x*-axis: core copy number (*i*_*o*_) present in population. Population sizes shown (in each bundle of three lines): *N* = 10*k* (upper), *N* = 2*k* (middle), *N* = 400 (lower). Duplication rate *d* = 2 *·*10^−3^, strength of selection *s* = 2 *·*10^−4^. 𝕍*/*𝔼-values calculated as averages across 100 replicates for each choice of *N* and *β*. Vertical line (*i*^*^ = 400) to ease comparison with empirical data.

## Bibliography

1. Hermann J. Muller. The relation of recombination to mutational advance. Mutation Research/Fundamental and Molecular Mechanisms of Mutagenesis, 1(1):2–9, 1964.

2. John Haigh. The accumulation of deleterious genes in a population - Muller’s ratchet. Theoretical Population Biology, 14(2):251–267, 1978.

3. Michael M. Desai. Haigh (1978) and Muller’s ratchet. Theoretical Population Biology, 133: 19–20, 2019.

4. Wolfgang Stephan, Lin Chao, and Joanne G. Smale. The advance of Muller’s ratchet in a haploid asexual population: Approximate solutions based on diffusion theory. Genetics Researc, 61(3):225–231, 1993.

5. Isabel Gordo and Brian Charlesworth. The degeneration of asexual haploid populations and the speed of Muller’s ratchet. Genetics, 154(3):1379–1387, 2000.

6. Richard A Neher and Boris I Shraiman. Fluctuations of fitness distributions and the rate of Muller’s ratchet. Genetics, 191(4):1283–1293, 2012.

7. Adrian González Casanova, Charline Smadi, and Anton Wakolbinger. Quasi-equilibria and click times for a variant of muller’s ratchet. Electron. J. Probab., 28:1–27, 2023.

8. Joseph Felsenstein. The evolutionary advantage of recombination. Genetics, 78(2):737–756, 1974.

9. John Maynard Smith. The evolution of sex. Cambridge University Press Cambridge, 1978.

10. Alexey S Kondrashov. Deleterious mutations and the evolution of sexual reproduction. Nature, 336:435–440, 1988.

11. Brian Charlesworth. Mutation-selection balance and the evolutionary advantage of sex and recombination. Genetics Research, 55(3):199–221, 1990.

12. Philip J Gerrish, Benjamin Galeota-Sprung, Fernando Cordero, Paul Sniegowski, Alexandre Colato, Nicholas Hengartner, Varun Vejalla, Julien Chevallier, and Bernard Ycart. Natural selection and the advantage of recombination. bioRxiv, 2021.

13. Sidhartha Goyal, Daniel J Balick, Elizabeth R Jerison, Richard A Neher, Boris I Shraiman, and Michael M Desai. Dynamic Mutation–Selection Balance as an Evolutionary Attractor. Genetics, 191(4):1309–1319, 2012.

14. Alexey S Kondrashov. Muller’s ratchet under epistatic selection. Genetics, 136(4):1469–1473, 1994.

15. Thomas Wiehe. Model dependency of error thresholds: the role of fitness functions and contrasts between the finite and infinite sites models. Genetics Research, 69(2):127–136, 1997.

16. Kavita Jain. Loss of least-loaded class in asexual populations due to drift and epistasis. Genetics, 179(4):2125–2134, 2008.

17. Manisha Brahmachary, Audrey Guilmatre, Javier Quilez, Dan Hasson, Christelle Borel, Peter Warburton, and Andrew J. Sharp. Digital genotyping of macrosatellites and multicopy genes reveals novel biological functions associated with copy number variation of large tandem repeats. PLoS Genetics, 10(6):e1004418, 2014.

18. Jaanus Suurväli, Colin J Garroway, and Pierre Boudinot. Recurrent expansions of b30. 2-associated immune receptor families in fish. Immunogenetics, pages 1–19, 2021.

19. Yannick Schäfer, Katja Palitzsch, Maria Leptin, Andrew R Whiteley, Thomas Wiehe, and Jaanus Suurväli. Copy number variation and population-specific immune genes in the model vertebrate zebrafish. eLife, 13:e98058, 2024.

20. John F. Crow and Motoo Kimura. An introduction to population genetics theory. Harper & Row, London, 1970.

21. Ugo Fano. Ionization yield of radiations. ii. the fluctuations of the number of ions. Phys. Rev., 72:26–29, 1947.

22. Andreas Leidenroth and Jane. E. Hewitt. A family history of dux4: phylogenetic analysis of duxa, b, c and duxbl reveals the ancestral duxgene. BMC Evol Biol, 10:364, 2010.

23. Moritz Otto, Yichen Zheng, Paul Grablowitz, and Thomas Wiehe. Detecting adaptive changes in gene copy number distribution accompanying the human out-of-africa expansion. Human Genome Variation, 11(37), 2024. doi: 10.1038/s41439-024-00293-w.

24. Feng Zhang, Wenlin Gu, Matthew. E. Hurles, and James. R. Lupski. Copy number variation in human health, disease, and evolution. Annual Review of Genomics and Human Genetics, 10:451–481, 2009.

25. Kondrashov Fjodor A. Gene duplication as a mechanism of genomic adaptation to a changing environment. Proc Biol Sci., 279:5048–57, 2012.

26. Michael Lynch and John S Conery. The evolutionary demography of duplicate genes. In Genome Evolution, pages 35–44. Springer, 2003.

27. Shozo Yokoyama and F Bernhard Radlwimmer. The “five-sites’ rule and the evolution of red and green color vision in mammals. Molecular Biology and Evolution, 15(5):560–567, 1998.

28. Yulin Gai, Ran Tian, Fangnan Liu, Yuan Mu, Lei Shan, David M Irwin, Yang Liu, Shixia Xu, and Guang Yang. Diversified mammalian visual adaptations to bright-or dim-light environments. Molecular Biology and Evolution, 40(4), 2023. doi: 10.1093/molbev/msad063.

29. Xiaopei Wang, Yongsen Ruan, Lingjie Zhang, Xiangnyu Chen, Zongkun Shi, Haiyu Wang, Bingjie Chen, Miles E Tracy, Chung-I Wu, and Haijun Wen. The paradox of extremely fast evolution driven by genetic drift in multi-copy gene systems. eLife, 13, 2024.

30. Robert E. Kass and Adrian E. Raftery. Bayes factors. Journal of the American Statistical Association, 90(430):773–795, 1995. doi: 10.1080/01621459.1995.10476572.

